# Pathogenic fungus expresses effector proteins in combination with a symbiotic virus to behaviourally manipulate housefly hosts

**DOI:** 10.1101/2024.08.20.608796

**Authors:** Sam Edwards, Knud N. Nielsen, Ian Will, Charissa de Bekker, Ioly Kotta-Loizou, Henrik H. De Fine Licht

**Affiliations:** Department of Plant and Environmental Sciences, University of Copenhagen, Frederiksberg C., 1871, Denmark; Living Systems Institute, Department of Biosciences, College of Life and Environmental Sciences, University of Exeter, Exeter, EX4 4QD, United Kingdom; Novo Nordisk Foundation Center for Protein Research, Faculty of Health and Medical Sciences, University of Copenhagen, Copenhagen N, 2200, Denmark; Department of Biology, University of Central Florida, Orlando, 32816, USA; Department of Biology, Microbiology, Utrecht University, Utrecht, 3584, The Netherlands; Department of Life Sciences, Faculty of Natural Sciences, Imperial College London, London, SW7 2AZ, United Kingdom; Department of Clinical, Pharmaceutical and Biological Science, School of Life and Medical Sciences, University of Hertfordshire, Hatfield, AL10 9AB, United Kingdom

## Abstract

Host manipulation by pathogens and parasites is a widespread phenomenon, but the molecular mechanisms are poorly understood. We investigated the summiting disease caused by the fungus *Entomophthora muscae* in houseflies, where infected flies climb to elevated positions and die, releasing infectious conidia. We performed dual-RNA sequencing of fly heads at different time points and identified candidate genes from both the host and the pathogen that may be involved in this summiting phenotype. Surprisingly, we also detected a high abundance of a novel positive sense single-stranded (+ss-RNA iflavirus in infected fly heads. We show that the virus load increases over time and shows signs of accumulation in fly heads and thoraces. We also reveal predicted interactions between fungal secreted proteins and insect host proteins related to neurological and immune functions, suggesting a possible role of these proteins in host manipulation. Furthermore, we find that *E. muscae* encodes a homologue of ecdysteroid UDP-glucosyltransferase (*egt*), a gene that has been implicated in host manipulation by other pathogens. Our study reveals a complex interplay between a fungus, a virus and a fly, and suggests that convergent evolution of *egt* for host manipulation mechanisms may occur in different pathogens.

## Background

Host specialisation has led to the evolution of the ability of pathogens to manipulate their hosts’ behaviour to increase transmission; a term coined the extended phenotype (Dawkins, 1982; Poulin & Maure, 2015). Striking pathogen host manipulations are mostly observed in invertebrates, however there are example of behavioural manipulation in vertebrates (Lafferty & Shaw, 2013). Within pathogen – invertebrate host manipulations, we see multiple strategies, including increased exposure for predation (such as surface swimming by amphipods to increase predation by birds (Bauer *et al*., 2005)). Equally striking manipulated host behaviours include parasitoid pupae protection in ladybirds (Maure *et al*., 2011) and spiders (Takasuka *et al*., 2015) or alteration in microhabitat choice, such as water-seeking in nematomorph-infected crickets (Ponton *et al*., 2011) and praying mantids (Obayashi *et al*., 2021) for parasite reproduction, or elevation seeking in fungal pathogen infected ants (Hughes *et al*., 2011) and flies (Evans, 1989) for increased infectious propagule transmission.

The manipulated behaviour whereby the host is forced to seek elevated positions to increase pathogen transmission is coined summiting disease (or tree top disease or ‘*Wipfelkrankheit’*). Summiting disease in a lepidopteran-baculovirus system has been found to be linked to viral hijacking of the expression of three host genes involved in visual perception, thus manipulating the insect to phototactically climb (Liu *et al*., 2022). Furthermore, a viral gene horizontally acquired from the insect called ecdysteroid UDP-glucosyltransferase (*egt*) and preventing molting in larvae, was found to be involved in eliciting climbing behaviour in the insect host (Hoover *et al*., 2011; Han *et al*., 2015). Lepidopteran larvae are hyperactive prior to death, a behaviour found to be regulated by the viral protein tyrosine phosphatase (*ptp*) (van Houte *et al*., 2013). However, the role of both *egt* and *ptp* depend on the lepidopteran-baculovirus system, as *ptp* has been shown to play a role in brain tissue infection rather than behaviour (Katsuma *et al*., 2012), and has no effect on hyperactivity (Kokusho & Katsuma, 2021) or summiting behaviour (Ros *et al*., 2015).

Behavioural manipulation of hosts infected with fungal pathogens has evolved multiple times within the fungal orders Hypocreales and Entomophthorales, but little is known about the underlying molecular mechanisms (Shang *et al*., 2015; Lovett *et al*., 2020; de Bekker *et al*., 2021). Pairwise comparisons using dual-RNA sequencing of *Ophiocordyceps* fungus-infected ants has revealed genes, pathways and mycotoxins potentially involved in behavioural manipulation, with a particular focus on summiting behaviour (de Bekker *et al*., 2015; Will *et al*., 2020). Although a complex phenotype like pathogen behavioural manipulation is likely polygenic and may involve multiple metabolic mechanisms working in concert (Herbison, 2017; de Bekker *et al*., 2021), advances are being made in testing candidate genes identified during gene expression analyses, such as an *Ophiocordyceps* mycotoxin related to aflatrem and linked to staggering behaviours seen prior to summiting (Beckerson *et al*., 2023). The convergent evolution of summiting disease multiple times across two fungal orders and a viral clade suggests that these taxonomically diverse pathogens target conserved insect phototactic metabolic traits and may use similar underlying mechanisms to manipulate host behaviours.

The obligate insect fungal pathogen *Entomophthora muscae* provides a tractable system for observing and defining specific manipulated behaviours (De Fine Licht *et al*., 2023). Behavioural manipulation by *E. muscae* occurs six or seven days post infection in houseflies (*Musca domestica*). Within six hours before sunset, the housefly host is forced to climb to an elevated position, glue its mouthparts to the substrate surface and terminate with the moribund zombie fly being prostrate (head down and abdomen elevated) with raised wings for optimal conidia dispersion from the abdomen (Krasnoff *et al*., 1995; Elya *et al*., 2018, 2023; Elya & De Fine Licht, 2021; De Fine Licht *et al*., 2023). To uncover the molecular mechanisms of these host moribund behaviours, here we first created a detailed ethogram of the manipulated behaviours in the *E. muscae*-housefly system to select the optimal time points for sample collection for dual-RNA sequencing. Second, differential gene expression analysis of pooled male housefly heads identified candidate host and fungal genes involved in behavioural manipulation, but also found high numbers of reads mapping to an iflavirus. Third, putative fungal small secreted proteins (SSPs) were analysed for predicted host-fungus protein-protein interactions (PPI) against the housefly proteome and the virus.

Here we show that PPI analysis predicted no interactions between six candidate SSPs and virus putative open reading frames (ORFs), but interactions were found with host proteins linked to neurological functions and disorders, suggesting that the fungal SSPs likely only bind to host proteins and not the virus. However, we find that the main component of total RNA from infected fly heads belongs to the iflavirus, a positive sense single-stranded (+ss) RNA virus. This virus has been identified previously as associated with *E. muscae* (Coyle *et al*., 2018), and our investigations into this virus confirms its taxonomic position amongst the *Iflaviridae* family, but highlight phylogenetic divergence of this group from insect-infecting viruses. Furthermore, PPI analyses of virus putative ORFs reveal binding between one viral protease with insect host proteins involved in kinesin motor domains and host cell nuclei, suggesting the virus may replicate within the host. Finally, using reverse transcription and polymerase chain reaction (RT-PCR) over the course of the infection of a new fungal isolate containing two viruses, we find viral load increases over time systemically with a tendency to concentrate in the fly head and thorax, where the major components of the insect central nervous system are located. We suspect that the build-up of this fungus-associated virus in the head and thorax of the fly, and the small number of fungal RNA reads identified using RNAseq of fly heads, suggests that the symbiotic virus may play a role in the behavioural manipulation of the *E. muscae*-housefly system.

## Results

### The *E. muscae* behavioural manipulation programme

The stereotypical *Entomophthora muscae* moribund manipulated behaviours of the host have been characterised by other studies in houseflies (Krasnoff *et al*., 1995) and *Drosophila melanogaster* (Elya *et al*., 2018, 2023). In our study system, the stereotypical behaviours of summiting, proboscis extension and wing raising were confirmed (Figure 1). However, within these three behaviours, we characterised specific sub-behaviours to better standardise specimen sampling for dual RNA sequencing due to the natural variability in the timing and duration of these three main behaviours. By creating a detailed ethogram for this system (Figure 1; Supplementary Figure 1; Supplementary Methods), we find that these different behaviours happen within six hours of subjected dark (lights-off) and this sequence of manipulated behaviours lasts on average *ca.* two hours (130±9.5 minutes) from start to finish (Figure 1; Supplementary Figure S1). In fact, live observations of the zombie houseflies show that on average, summiting (SM), proboscis extension (PB) and wing raising (WG) last respectively 78±12.6, 43±6.8 and 9±2 minutes in this system. Summiting behaviour in houseflies was predominantly observed six days post infection, however not all flies died after six days (Supplementary Figure S1). For surviving flies, the observed behaviours were aberrant as compared to uninfected flies, demonstrating that *E. muscae* infection also affects the normal behaviour of houseflies (see ‘Infected (2)’ in Supplementary Figure S1). The variation in time of death, being on day 6 or day 7, could be due to either the nature of the pathogen, as *E. muscae* conidia germination takes 2-24 hours and thus the day 7 manipulation may be due to delayed germination and entry of the fungus into the fly’s haemolymph (Brobyn & Wilding, 1983), or individual variation in host response to infection.

**FIGURE 1.**
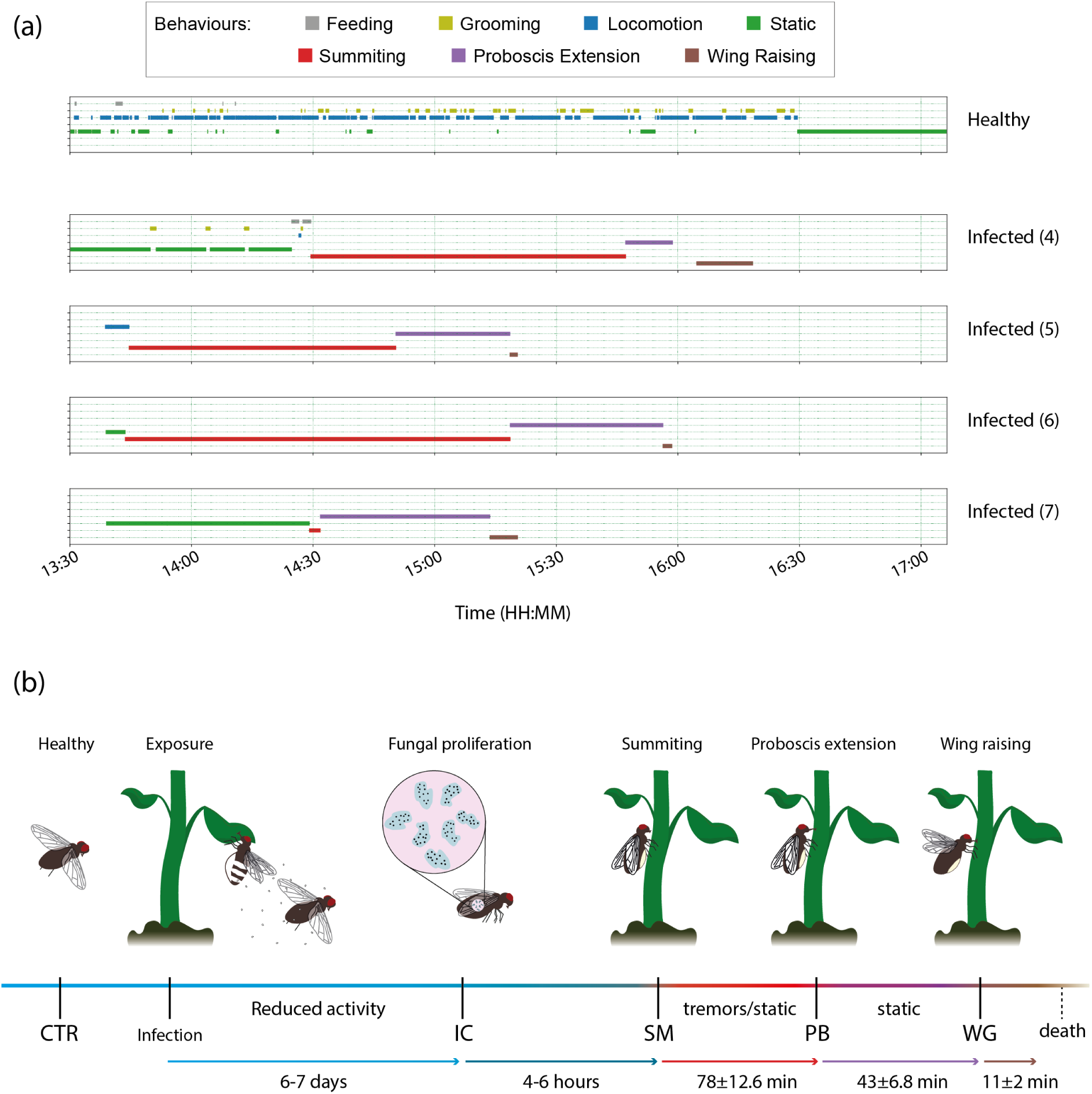
Timeline of the manipulated behaviours of live observed infected houseflies six days post infection. Houseflies were live observed and their behaviours recorded from 13:30 to 17:05, where the artificial lighting in the lab would turn off at 18:00. **(a)** Timeline of the observed behaviours, the start time and duration of behaviours vary for each individual observed, requiring identification of sub-behaviour (not represented here) to standardise specimen sampling. **(b)** Summarised timeline of an *E. muscae* infection and the associated observed manipulated behaviours. Points at which samples were collected for RNAseq analyses are indicated by vertical lines along the timeline.

### Housefly molecular response to manipulation

To identify candidate host genes potentially involved in the manipulated behaviours observed above, we sequenced the transcriptome of housefly heads collected during the SM, PB, and WG phenotypic time points. As controls, we also included infected flies (IC) collected at six days post infection, corresponding to 4-6 hours before the onset of the behavioural manipulation, and uninfected control flies (CTR) of the same age. For each treatment, the mean number of sequenced reads was between 22.8 and 24.4 million (Supplementary Table S1). A mean of 89% ±2 mapped to the fly host genome in uninfected control (CTR) flies. Read mapping rate in the infected groups were as expected lower and decreased with disease progression and build-up of fungal biomass (Hansen & De Fine Licht, 2017), with an average of 25, 21, 14 and 10% reads mapping to the host genome for infected control (IC), SM, PB and WG, respectively (Supplementary Table S1). In total, 14,550 out of 16,135 genes in the house fly genome had at least one read mapping.

Principal component analyses (PCA) revealed that the five different treatments clustered separately, with the exception of PB and WG, which have 104 genes that are differentially expressed in pairwise comparisons (Figure 2a,c,g,h). In fact, pairwise comparisons with uninfected flies revealed that over the course of infection, there were increasing numbers of differentially expressed genes (DEGs) in infected flies, from 1779 between CTR and IC, and 4872 between CTR and WG, the majority of which were downregulated (Figure 2c-h). To investigate general functions of housefly DEGs, we performed gene set enrichment analyses using gene ontology (GO) terms. These analyses were performed across a total of 231 gene sets, comparing each behavioural phenotype to the previous one during disease progression to gain insight into the gene set changes over time. We did this through comparing IC to CTR, SM to IC, PB to SM, and WG to PB. Similarly to DEGs, there is a negative trend in enrichments over time in host gene sets across infection time (Supplementary Figure S1-4).

**FIGURE 2.**
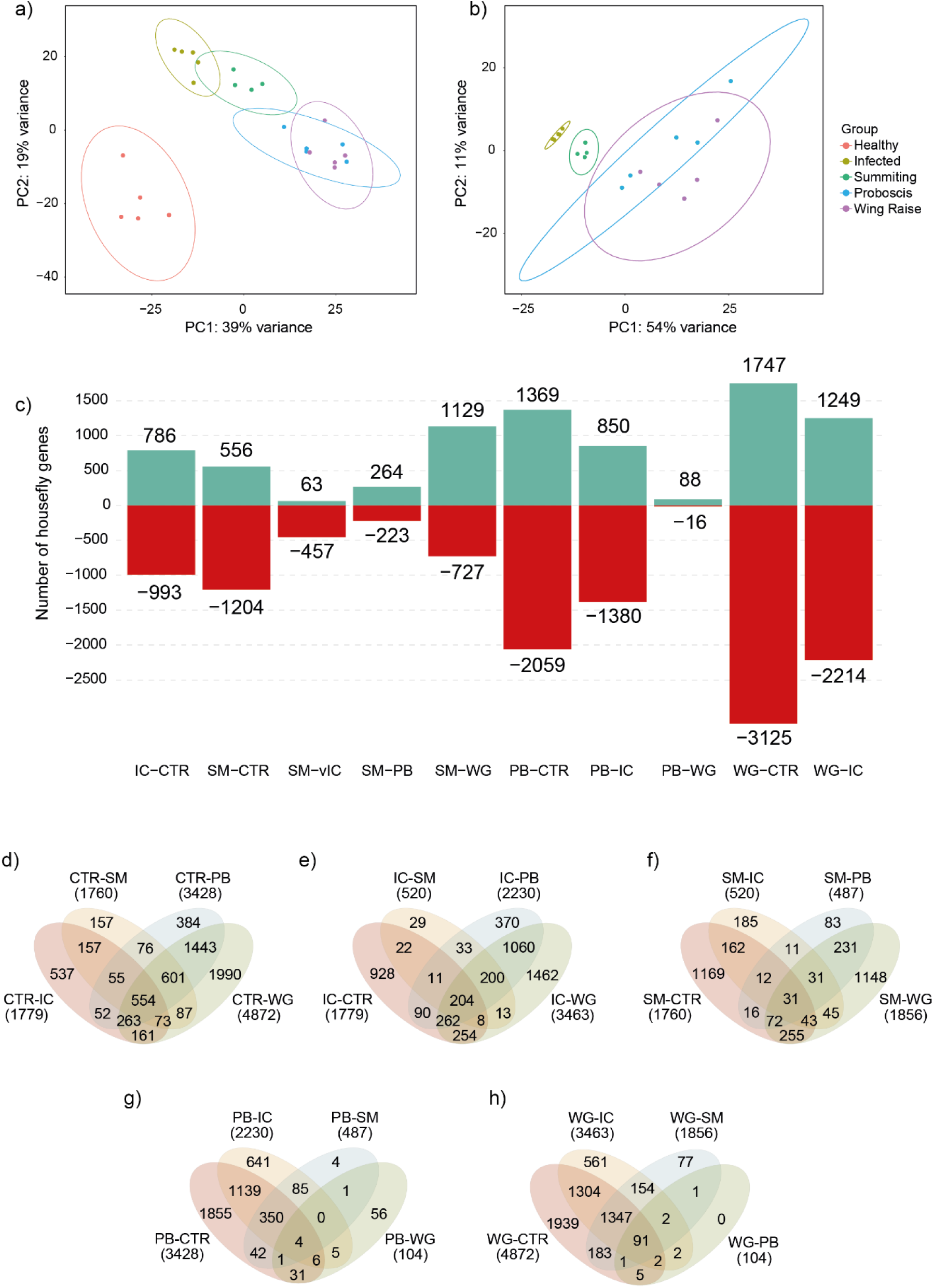
Differentially expressed genes (DEGs) of genes of *Musca domestica* during manipulation by *Entomophthora muscae*. **(a)** Principal component analysis for RNAseq rlog transformed counts of genes from houseflies and **(b)** from *E. muscae* samples. **(c)** Pairwise comparisons of DEGs across infected time points. **(d)** Venn diagram analysis of the pairwise comparisons identified for shared and unique DEGs of *E. muscae*-infected houseflies for control, **(e)** infected control, **(f)** summiting, **(g)** proboscis extension, and **(h)** wing raising flies.

Gene sets enriched in the comparison between IC and CTR are assumed to be linked to infection by *E. muscae* and not to be linked with behavioural manipulation. In total, IC had 20 and 10 gene sets respectively upregulated and downregulated as compared with CTR with false discovery rate below 5% (Supplementary Figure S1). Enrichment in gene sets upregulated in IC involved immune functions (defence response to bacterium; innate immune response; regulation of Rho protein signal transduction) and response to stress (Supplementary Figure S1). Significantly downregulated gene sets in IC compared to CTR included steroid hormone mediated signalling pathway, odorant binding, and nuclear steroid receptor activity (Supplementary Figure S1).

During summiting behaviour, we saw 12 upregulated and 50 downregulated gene sets as compared with IC (Supplementary Figure S2). With the exceptions of peptidoglycan catabolic process and innate immune response that were not enriched and biological processes that remained downregulated in the pairwise comparison of SM and IC, all other gene sets enriched in the IC-CTR comparison are oppositely regulated (up in IC-CTR and down in SM-IC, and vice versa) (Supplementary Figure S1,2). The same pattern is seen in the gene sets involved in molecular processes and cellular components (Supplementary Figure S1,2), which to some extent likely reflect the difference in *E. muscae* disease progression from IC over SM to PB and WG. GO terms linked to ion (monoatomic, potassium) and ion transmembrane (calcium) transport, steroid hormone mediated signalling pathway, cell junction, and postsynaptic membrane were upregulated in SM (Supplementary Figure S2). Enrichment of these GO terms may suggest increased sensory activity and signal transmission during summiting behaviour (Supplementary Figure S2).

In total, we discovered seven upregulated and 23 downregulated gene sets during pairwise comparisons of PB and SM (Supplementary Figure S3). Immune defence responses, and intracellular protein transport molecular functions decrease further in PB. However, upregulation of sensory perception of smell, notch signalling pathway and cyclic nucleotide biosynthetic process can be seen, all of which were not enriched in prior pairwise comparisons. Molecular functions relating to structural constituent of the cuticle, olfactory receptor activity and phosphorus-oxygen lyase activity were upregulated. All gene sets upregulated during SM and IC comparisons were not differentially enriched between PB and SM (Supplementary Figure S2,3).

Pairwise comparisons of WG and PB revealed 10 downregulated gene sets and no significant upregulated gene sets (Supplementary Figure S4). Gene sets involved in molecular functions cell redox homeostasis, translation, tricarboxylic acid cycle and proton motive force-driven ATP synthesis were downregulated. Sets involved in biological processes structural constituent of ribosome and cytochrome-c oxidase activity were also differentially downregulated. Cellular components for mitochondrion, mitochondrial inner membrane and ribosome were also downregulated in WG. Downregulation in all gene sets likely indicates that the host houseflies were dying during WG (Supplementary Figure S4).

### Genes to force your host to su(b)mmit

PCA of fungal DEGs revealed that the four different treatments clustered separately, with the exception of PB and WG, which share one gene that is differentially expressed in pairwise comparisons (Figure 2b). The lack of differences in fungal DEGs between PB and WG suggests that the WG phenotype may be an artefact of fungal growth and changes in pressure within the host as hypothesised by other *E. muscae* studies (Figure 3a,d,e) (Elya & De Fine Licht, 2021). Thus, assuming that genes involved in fungal behavioural manipulation would only be active during summiting, we aimed to identify fungal genes that are only differentially expressed during summiting.

**FIGURE 3.**
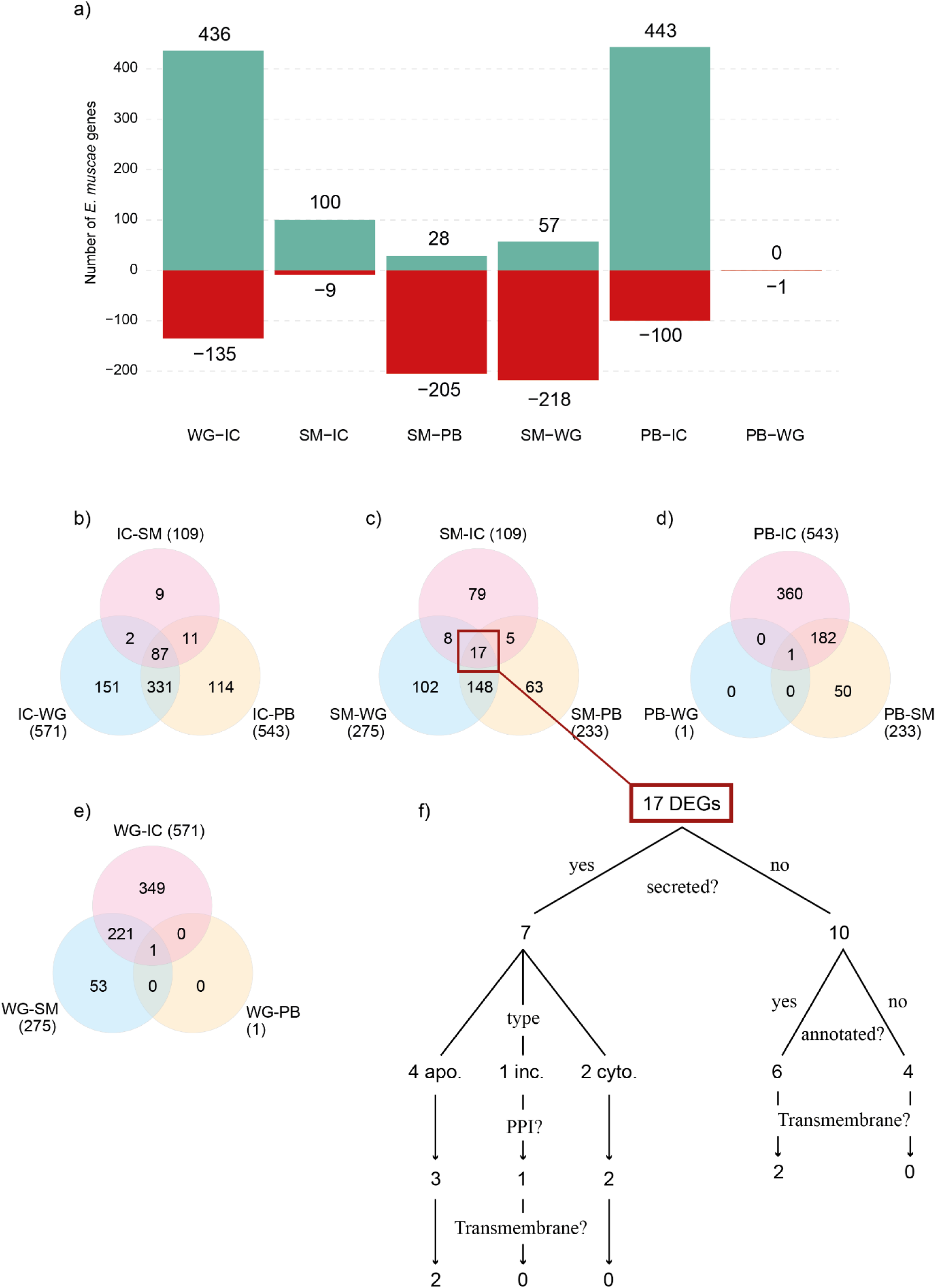
Differentially expressed genes (DEGs) of *Entomophthora muscae* during manipulation of infected male *Musca domestica*. **(a)** Pairwise comparisons of DEGs across infected time points. Number of genes in green are up-regulated and red are down-regulated with the exact number written above and below each bare, respectively. **(b)** Venn diagram analysis of the pairwise comparisons identified for shared and unique DEGs of *E. muscae* for infected control, **(c)** summiting, **(d)** proboscis extension, and **(e)** wing raising flies. **(f)** A flow diagram illustrating the known characteristics of the 17 genes uniquely expressed during summiting behaviour identified during DGE analysis.

Pairwise comparisons of summiting behaviour DEGs to the previous (IC) and subsequent (PB and WG) treatment time points identified 17 DEGs (Figure 3b). Out of these 17 genes, 13 were increasingly upregulated over time, one decreasing, and three were significantly upregulated at the summiting point, but downregulated at IC PB and WG (Figure 4). Of these 17 DEGs, seven had a SignalP domain (out of 1792 in total with a SignalP domain in the *E. muscae* transcriptome) and were predicted by the EffectorP software to be small secreted peptides (SSPs), of which two were cytoplasmic, four apoplastic and one inconclusive as the predicted type was both apoplastic and not apoplastic (using ApoplastP) (Figure 3f & 4a-g). As expected, none of the SSPs could be identified to have known molecular functions, although HBVY01021106 (apoplastic effector) is predicted to have a transmembrane protein (Figure 4c). Looking at the expression profiles of these SSPs over the course of manipulation, we see that HBVY01016605 and HBVY01021106 (Figure 4b,c) are upregulated uniquely during summiting behaviour, which indicates a likely involvement in the summiting phenotype of *E. muscae* infected houseflies.

**FIGURE 4.**
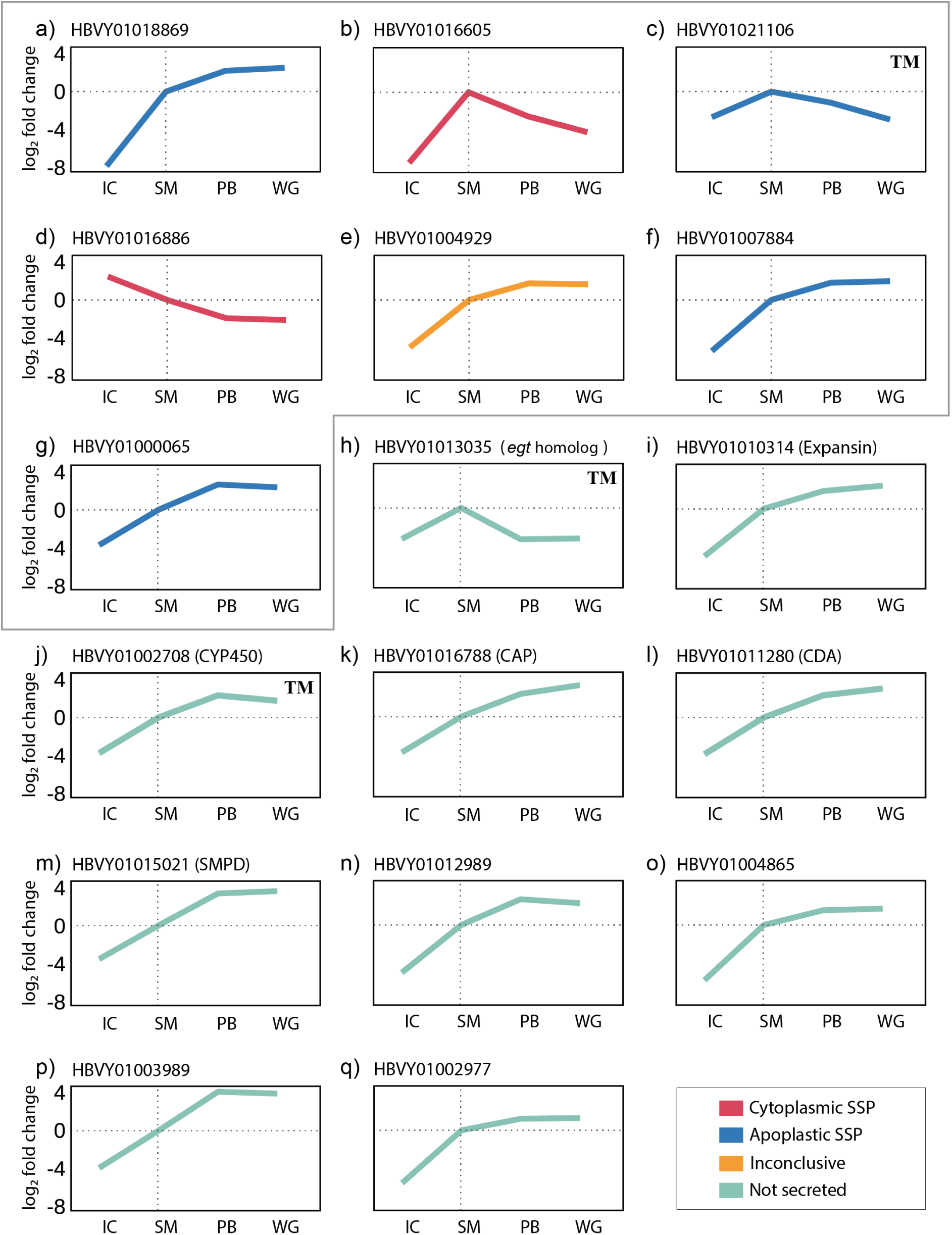
Expressions profiles of 17 *E. muscae* DGE identified through pairwise comparison of summiting behaviour versus all other treatments (figure 3f). **(a)** – **(q)** In each panel, the x-axis shows the four behaviours from infected control (IC), summiting (SM), proboscis extensions (PB), and wing raising (WG), and y-axis show the log_2_ fold-change in expression. Line colours depict the predicted type of effector, or whether they are non-secreted genes, with red: Cytoplasmic SSP, blue: Apoplastic SSP, orange: Inconclusive as either apoplastic or cytoplasmic, green: Not secreted. Genes identified as SSPs are encircled in one large grey box. TM in the top right-hand corner of the plots indicate which genes are predicted to have transmembrane domains. Gene abbreviations: CYP450 = Cytochrome P450 monooxygenase; CAP = Catabolite activator protein; CDA = Chitin deacetylase; SMPD = Sphingomyelin phosphodiesterase.

Of the ten non-SSPs identified to be differentially expressed in summiting behaviour, all but one (HBVY01013035; Figure 4h) have increased expression over time (from IC to WG) (Figure 4i-q). Six of these DEGs could be tentatively functionally annotated using homology searches (Figure 4h-m). HBVY01002708 encodes a homologue of cytochrome P450 monooxygenase (CYP450) FUS8 and a transmembrane protein (Figure 4k). In *Fusarium verticillioides*, a common maize fungal pathogen, the P450 FUS8 gene is proposed to belong to a fusarin biosynthetic gene cluster (Brown *et al*., 2012). Fusarins are mycotoxins, but *E. muscae* is not known to produce toxins (Boomsma *et al*., 2014; Elya & De Fine Licht, 2021). However it has been suggested that *E. muscae* is able to synthesise certain secondary toxins (De Fine Licht *et al*., 2017), and the presence of a CYP450 FUS8 homologue gene may suggest that *E. muscae* has genetic potential to produce fusarin-like metabolites. HBVY01010314 encodes for expansin (PF00967 - Barwin family) (Figure 4i), a protein involved in defense response to bacterium and fungi in plants. However, the function of expansins in fungi remains unknown (Narváez-Barragán *et al*., 2020). But expansins in phytopathogenic fungi may function as virulence factors or act in cell wall extension similar to expansins in plants (Chase *et al*., 2020). The gene HBVY01011280 encodes chitin deacetylase (CDA) (UniProtKB:Q06702) (Figure 4l), an enzyme normally involved in synthesis and modulation of fungal cell walls. *E. muscae* is known to propagate over the week-long infection as wall-less protoplast cells to evade host immune detection (Boomsma *et al*., 2014), but has been found to develop a cell wall toward the end of the within-host life cycle. HBVY01016788 encodes a PFAM domain for cysteine-rich secretory protein family (PF00188), a subgroup of catabolite activator protein (CAP) superfamily, which is a large superfamily of pathogenesis-related proteins when present in pathogens (Figure 4k). The HBVY01015021 gene encodes sphingomyelin phosphodiesterase, an enzyme involved in the sphingolipid metabolic pathway. In the pathogenic fungus *Candida albicans*, sphingolipids are key for the normal growth of hyphae (Martin & Konopka, 2004). In *Cryptococcus neoformans*, these lipids play important roles in fungal pathogenicity (Singh & Del Poeta, 2011). Moreover, an increase in secondary metabolites resulting from the sphingolipid metabolism may be one of mechanisms used by the predatory fungus *Duddingtonia flagrans* to capture and kill nematodes (Liang *et al*., 2019).

Most notable however, is HBVY01013035 (Figure 4h), which encodes a PFAM domain for UDP-glucosyl transferase (PF00201) and a transmembrane protein. Ecdysteroid UDP-glucosyltransferase (*egt*) is a UDP-glycosyl transferase homologue and is a key gene responsible for inducing climbing behaviour in behaviourally manipulating baculoviruses (Hoover *et al*., 2011; Han *et al*., 2015). In *E. muscae*, this putative fungal *egt* gene is uniquely upregulated (ca. 4-fold up from IC and then down to PB) during summiting in *E. muscae*-infected flies (Figure 4h), indicating likely involvement in this manipulation phenotype. Phylogenetic analysis suggests that the nucleopolyhedrovirus (NPV) *egt* is divergent from those known in fungi, but also from the identified UDP *E. muscae* protein (Figure 5a). *E. muscae* UDP had 22-29% sequence similarity with the top 10 *Baculoviridae* hits with the lowest e-value in BLASTp. As *egt* is known to add a sugar moiety to 20E in the insect molting hormone biosynthesis pathway, thus disabling 20E’s function (O’Reilly *et al*., 1992), we looked into the expression profiles of all known host genes involved the hormone biosynthesis pathway. The genes encoding for ecdysteroid 25- and 2-hydroxylase (*phm* and *sad*) are significantly downregulated specifically during summiting behaviour. The enzyme ecdysteroid 20-monooxygenase/CYP314A1/Shade (*Shd*), which converts ecdysone into 20E, decreases in expression during PB, correlating with an increased expression of 26-hydroxylase/CYP18A1, the enzyme encoded to inactivate the active hormone 20E (Figure 5b; Supplementary Figure 5) (Guittard *et al*., 2011; Niwa & Niwa, 2014).

**FIGURE 5.**
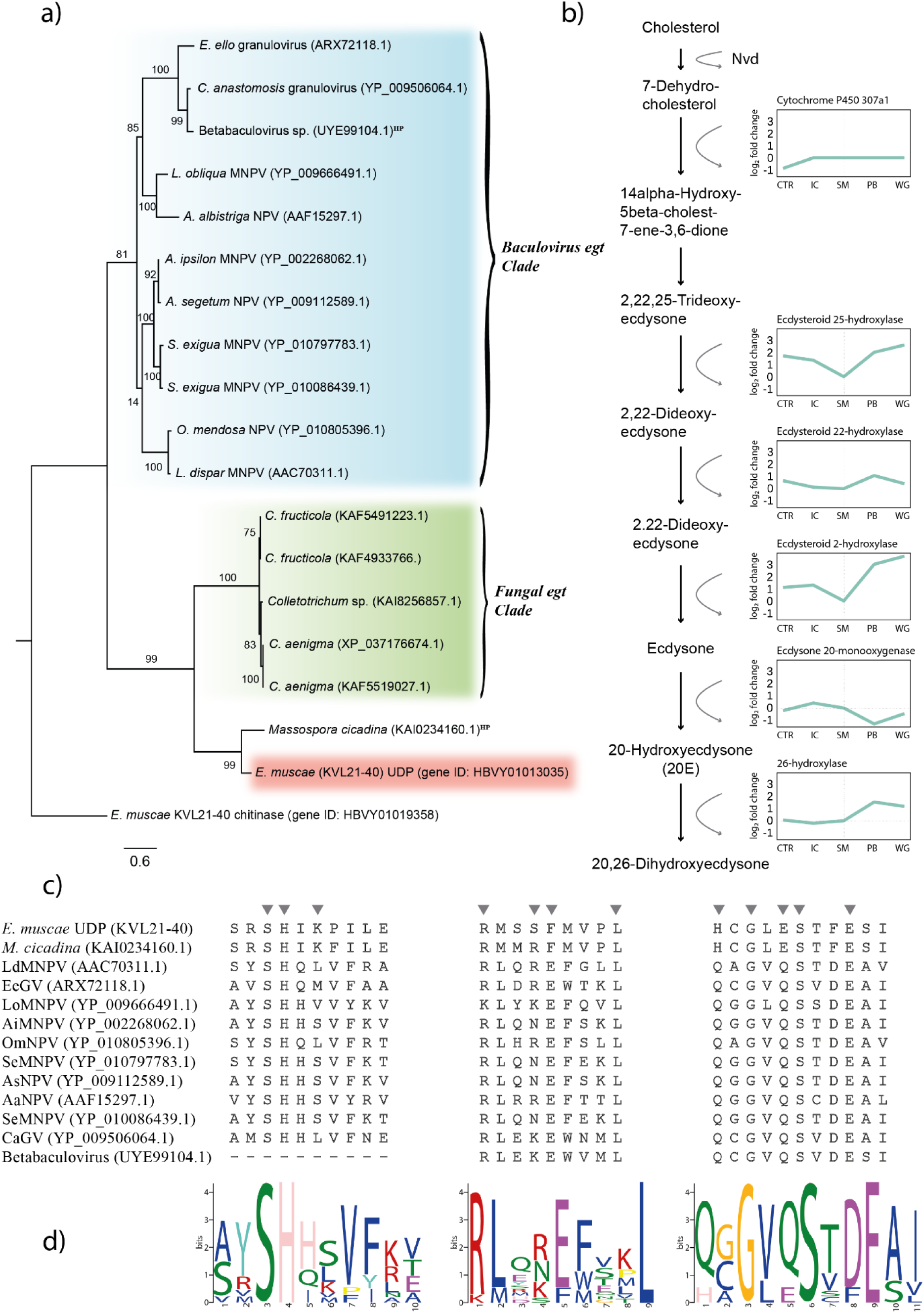
*E. muscae egt* homologue may be involved in summiting disease in houseflies. **(a)** Maximum likelihood phylogenetic analysis of the UDP-glucoronosyl and UDP-glucosyl transferase (UDP), and ecdysteroid UDP-glucosyltransferase (*egt*) genes from fungi and baculoviruses based on amino acid sequences. Bootstrap percentages are shown for each node, and an *E. muscae* chitinase was used as outgroup. Blue, orange and green highlights depict clades of known UDP genes, red highlight shows the upregulated *E. muscae* gene identified through DGE analysis. Abbreviations: MNPV = Multiple nucleopolyhedrovirus; NPV = Nucleopolyhedrovirus; HP = Hypothetical Protein. **(b)** Transcriptional profiling of the housefly molting hormone biosynthesis pathway from cholesterol to the 20-Hydroxyecdysone (20E) molting hormone that are identified to be involved in some baculovirus-caterpillar systems. Expression plots similar to figure 4 for each housefly enzyme in the pathway is shown. Missing enzymes have never been identified in house fly genomes. **(c)** Amino acid alignment of the active site residues (12 out of 14 displayed) for the GT1_Gtf-like domain identified among the candidate *E. muscae* UDP gene. The triangles above the sequences represent the location of the active sites. **(d)** Sequence motifs of the active site residues for each active site group.

In order to verify if the putative *E. muscae egt* contained similar active site residues as the lepidopteran larvae manipulating baculoviruses we mined the conserved domain database (Wang *et al*., 2023), which revealed that the putative *E. muscae egt* contains homologous amino-acid residues at 14 out of 14 active binding sites (12 of which are represented in Figure 5c), sharing 10-50% sequences similarity with selected *Baculoviridae egt* genes (Figure 5c). The *E. muscae egt* active site residues resemble those of the baculoviral motifs (Figure 5d), although any effect of the *E. muscae egt* gene may have on the host’s molting hormone pathway remains inconclusive (Supplementary Figure S5).

### Overrepresentation of viral RNA reads in infected housefly heads

RNAseq mapping in *E. muscae*-infected houseflies revealed that ∼25% of reads mapped to the housefly genome and ∼1% to the *E. muscae* transcriptome (Figure 6a). The majority of the remaining reads mapped to the genome of the *Entomophthora* virus (EV) from the housefly-isolated *E. muscae* genotype (KVL21-40) which was used for infections in specimens for RNAseq (Figure 6a; Supplementary Table S2). This virus is a member of the family *Iflaviridae* (Type I), with a genome length of ∼8.7 kbp and encompasses two large partially overlapping open reading frames (ORF) (Coyle *et al*., 2018). The 5’-proximal 1900 amino acid (AA) polyprotein encodes three structural capsid proteins and helicase, and the 3’-proximal non-structural 1013 AA ORF encodes the RNA-dependent RNA polymerase (RdRp) and 3C protease (Figure 6b).

**FIGURE 6.**
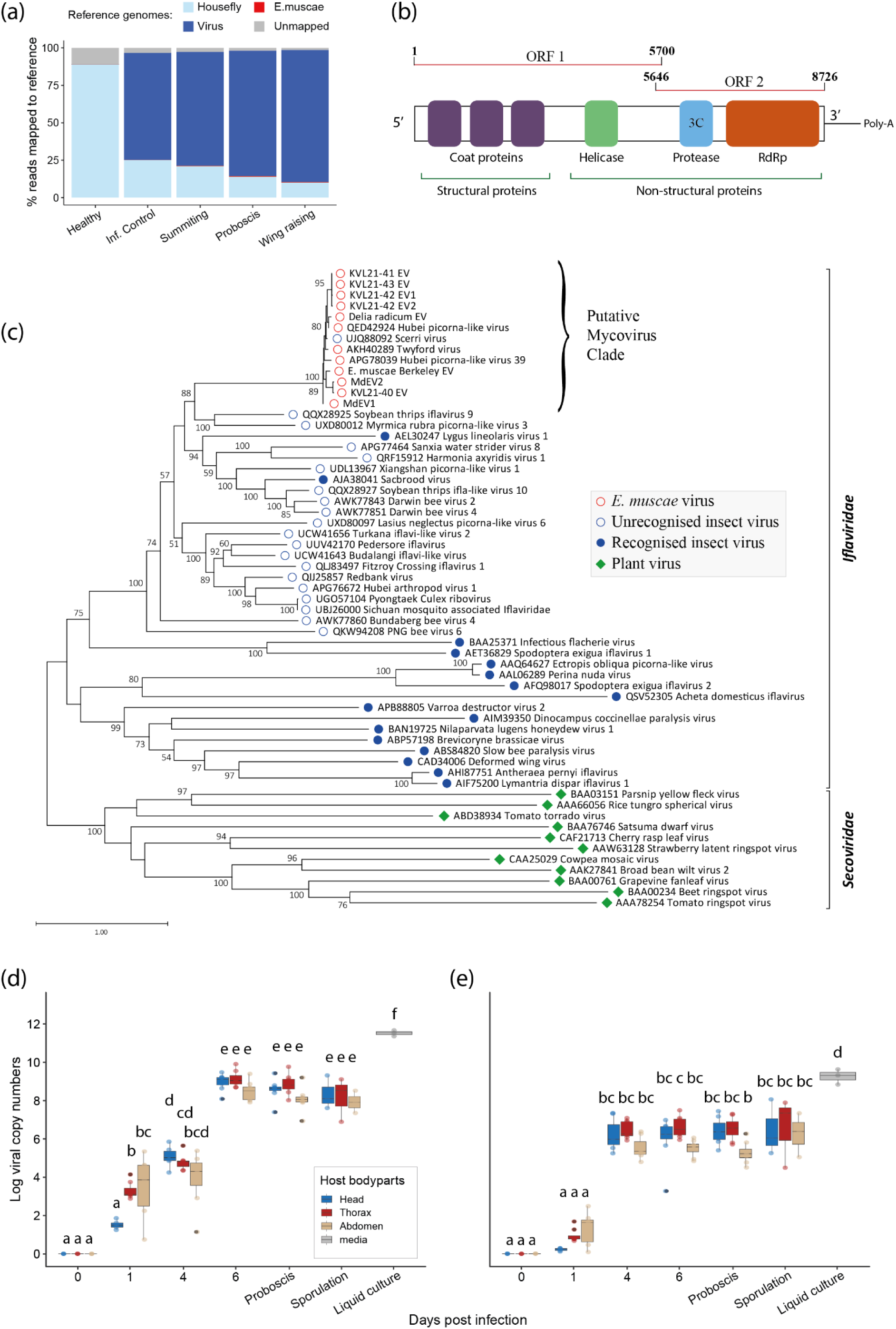
Symbiotic viruses are involved in an *Entomophthora muscae* infection of houseflies. **(a)** Percentage of RNAseq reads mapping to either the housefly, *E. muscae,* or virus (EV) genome over the course of disease progression time points. **(b)** Schematic diagram of the predicted 8726 base-pair EV genome structure. Numbers on either end of the ORFs indicate nucleotide positions. Boxes indicate the position of characterised proteins. **(c)** A maximum likelihood phylogenetic analysis of the RNA-dependent RNA polymerase (RdRP) amino-acid sequences of newly sequenced mycoviruses KVL21-41 EV (*Drosophila* isolated), KVL21-43 EV (*Drosophila* isolated), KVL21-42 EV1 and EV2 (*Drosophila* isolated), *E. muscae* Berkeley EV (*Drosophila* isolated), *Delia radicum* isolated *E. muscae* EV, KVL21-40 EV, MdEV1 and MdEV2, together with related viruses. Bootstrap percentages (100 replicates) are shown. Members of the *Secoviridae* family was used as outgroup. Tips labelled with circles indicate *Iflaviridae* and rhombuses *Secoviridae*; solid shapes, in contrast to outlines, indicate viruses from officially recognised species; colours indicate different virus hosts, red for fungi, blue for arthropods and green for plants. **(d)** Viral copy numbers in the head, thorax and abdomen of houseflies infected with *E. muscae* during the course of infection for MdEv1 and **(e)** MdEv2. Each coloured dot in the plot represent the virus titre of the body part of a single fly. Different letters indicate statistically significant differences.

Due to the overrepresentation of viral reads in infected housefly heads, investigation into the role of the virus was prioritised. The newly isolated *E. muscae* fungus genotype (KVL22-01) revealed the presence of two distinct EVs, MdEV1 and MdEV2, sharing 82.3% pairwise identity in the RdRp region. Both viruses were confirmed as members of the *Iflaviridae* through BLAST searches of their RdRp region sequences. Phylogenetic analyses of these two new EVs confirm their clustering with the KVL21-40 EV found in high numbers in housefly heads in this study, and with other known EVs, i.e. Twyford and Hubei picorna-like virus 39 from insect samples, viruses directly sequenced from *E. muscae* isolate (Coyle et al. 2018; **Chapter 6**), and the Scerri virus that was sequenced from *Haemaphysalis bancrofti* ticks in Australia (Gofton *et al*., 2022). Phylogenetic analysis places the EVs in the *Iflaviridae*, a virus family known to be arthropod, but primarily insect infecting (Valles *et al*., 2017). The phylogenetic placement of EVs in the *Iflaviridae* is in accordance with previous studies (Coyle *et al*., 2018; Myers *et al*., 2020). However, by using RdRp sequences from 12 EVs, our phylogeny shows a divergence from recognised insect-infecting viruses (Figure 6c), supporting a possible horizontal transmission from insects to entomopathogenic fungi in this putative mycovirus clade as suggested previously (Coyle et al. 2018). Why the Scerri virus is placed within the EVs remains to be determined, but this finding may propose that the original tick specimen could have had a fungal infection.

With the finding that EVs are found in high numbers in housefly heads used for RNAseq, we quantified and assessed the distribution of MdEV1 and MdEV2 using RT-qPCR on different host parts of infected houseflies, *viz.* head, thorax and abdomen, over the course of an *E. muscae* within-host life cycle. Specifically, we assessed viral titre in uninfected (0 dpi), 1, 4 and 6 (morning of manipulation) dpi, during manipulation (proboscis extension), after *ca.* 12 hours of sporulation and in liquid fungal culture. Viral titres for MdEV1 and MdEV2 within the houseflies were significantly different (ANOVA: LR χ^2^_1_ = 13.486, P < 0.001), with MdEV1 being *ca.* 10-fold more abundant by the end of the within-host fungal life cycle with mean viral copy numbers of MdEV1 = 5876 (min = 1029, max = 17447) and MdEV1 = 593 (min = 90, max = 1717) per 20 ng total RNA.

There were significant differences in viral titres in host body parts, across dpi and in their interaction in both MdEV1 (ANOVA: LR χ^2^_2_ = 7.92, P = 0.019; ANOVA: LR χ^2^_5_ =1715.09, P < 0.001; ANOVA: LR χ^2^_10_ = 32.75, P < 0.001) and MdEV2 (ANOVA: LR χ^2^_2_ =15.82, P < 0.001; ANOVA: LR χ^2^_5_ = 1255.34, P < 0.001; ANOVA: LR χ^2^_10_ = 22.27, P = 0.014). At 1 dpi, MdEV1 is significantly higher in the thorax and abdomen of infected flies than in heads, viral titre in the latter being not statistically different to uninfected flies. By 4 dpi, there are higher numbers of MdEV1 in the head and the thorax than in the abdomen, however these differences are not statistically different. But viral titre in the head and thorax 4 dpi are statistically significantly higher than 1 dpi, but not in the abdomen. Viral titre significantly increases from 4 to 6 dpi, but is not significantly different thereafter in the host. However, the trend of higher mean copy numbers in heads and thorax than in the abdomen remains at 6 dpi, during PB manipulation and in sporulation, with the latter having slightly lower viral titre (Figure 6d). In MdEV2, there are no statistical differences between houseflies that are uninfected and 1 dpi, but there is more virus in the abdomen overall at 1 dpi. From 4 dpi, the viral titre of MdEV2 is statistically significantly higher from 1 dpi, but not from 4 dpi to sporulation, with the exception of viral titre in the thorax 6 dpi being significantly higher than in the abdomen of PB manipulated flies (Figure 6e). Both MdEV1 and MdEV2 show on average higher viral abundance in the head and the thorax than in the abdomen. Furthermore, we find significantly higher abundance of both EVs in liquid fungal cultures than in infected flies (Figure 6d,e), which suggests that MdEV1 and MdEV2 are transmitted with conidia during cadaver sporulation and replicate in fungal cells.

### Insect-fungus-virus protein-protein interactions

In summiting disease, pairwise comparisons identified 17 DEGs, seven of which were predicted to be effector proteins. Predicted protein-protein interactions (PPIs) of the seven fungal SSPs identified during DEG analysis showed that six fungal SSPs were predicted to interact with at least one host protein (Supplementary Table S3). The SSPs had mixed connectivity to fly proteins, ranging from one to 105 (median = 8.5). In total 125 of 18,827 fly proteins were predicted in PPI, where only one host protein was found to bind to one fungal protein, as opposed to one to four in the *Ophiocordyceps*-*Camponotus* system (Will *et al*., 2023). The translated protein from the HBVY01016605 gene that is differentially expressed during summiting binds to 105 host proteins. These host proteins are involved in transport and localisation GO processes in the lysosome, ribosome and oxidative phosphorylation pathways. Within these 105 host proteins, six genes encoded facilitated trehalose transporters (*Tret*) (four *Tret*1 and two *Tret*1-2 homologues) were predicted to have interactions. *Tret*1 allows for transmembrane transport of synthesised trehalose from the fat body to the haemolymph in *Polypedilum vanderplanki* flies (Kikawada *et al*., 2007). *E. muscae* interactions with trehalose synthesis may be implicated in the fungus’ strategy of outcompeting the host’s own trehalose metabolism (De Fine Licht *et al*., 2017). The protein encoded in HBVY01004929 and HBVY01007884 were predicted to interact with one host phosphoglycerate mutase 2 (PGAM2) and one formin-A gene, respectively. PGAM2 is involved in glucose metabolism and PGAM deficient *Drosophila melanogaster* embryos exhibit reduced muscle fusion and repair (Tixier *et al*., 2013), and in plants is involved in mitochondria-chloroplast tethering (Zhang *et al*., 2020). HBVY01018869 predictively binds to nine host proteins, eight encoding T-complex 1 protein and one encoding heat-shock protein 60A (HSP60), all of which are chaperonins and play cellular roles in protein folding. Of interest were two fungal genes, HBVY01021106 and HBVY01016886, which predictively interact with eight and 1 host proteins, respectively. The former gene is upregulated specifically during summiting (Figure 4b), whilst the latter is downregulated from IC to WG (Figure 4d), suggesting that HBVY01021106 may be involved in housefly summiting behaviour. These host genes encode for Down syndrome cell adhesion molecule-like protein (Dscam2), synaptotagmin 1, a partial tyrosine-protein phosphatase (*ptp*) 99A-like, and cytochrome b5. The host genes binding to the two SSPs have gene ontology (GO) processes such as chemotaxis, locomotion, axonogenesis, neurogenesis, neurotransmitter secretion and nervous system development. In particular, *ptp* binding is interesting due to its known involvement in host hyperactivity prior to summiting behaviour in baculovirus-caterpillar systems (Katsuma *et al*., 2012; van Houte *et al*., 2012; Kokusho & Katsuma, 2021).

For the virus-fungus PPI, 21 combinations were tested between the seven candidate fungal SSPs and three virus proteins (two ORFs and one partial protease), but no connectivity was found. This shows that the proteins secreted by *E. muscae* specifically during summiting do not interact with the virus, suggesting the summiting candidate SSPs only interact with the host.

We found seven PPI interactions out of the 56,481 EV-housefly PPI combinations tested, all seven host proteins predicted to bind were to a partial 40 amino acid sequence of the viral protease ORF, but no interactions were predicted with the entire two large ORFs (Supplementary Table S4). Three of these seven host transcripts are zinc finger, nuclear hormone receptor-types, but more specifically are an ecdysone receptor, a transcription HNF-4 homologue and a probable nuclear hormone receptor HR38, all with GO terms linked to host cell nuclei. The four other host transcripts involved in PPI are kinesin motor domains, which are involved in microtubule-based movement within host cells. Host microtubules are known to be directly involved in cytoplasmic viral transport (Döhner *et al*., 2005), suggesting that EVs may bind to host microtubules to be transported to and from host nuclei to replicate within insect cells.

## Discussion

In this study we find that *Entomophthora muscae* infected flies display stereotypical manipulated behaviours and identify candidate fungal genes that appear to influence summiting behaviour, notably a few specific fungal small secreted proteins (SSPs) and an *egt* homologue. In some baculovirus-lepidopteran systems, the horizontally acquired *egt* gene is responsible for inducing summiting disease in infected larvae (Hoover *et al*., 2011; Han *et al*., 2015). Although the function of the fungus *egt* homologue was not tested in this study, the systemic presence of a fungus-infecting insect virus leaves open speculation as to whom is behind the insect manipulation, or at least as to what role the virus plays in this three-way interaction. We find that the virus proliferates throughout the insect host during infection, with viral titres being slightly higher in the head and thorax compared to the abdomen. Furthermore, low fungal and high viral RNA read numbers in housefly heads, and high viral titres in liquid culture of the fungus indicate that the virus must replicate within both insect and fungal cells. However, protein-protein interactions analysis shows that a partial sequence from the viral protease binds to domains involved in host cytoplasmic transport, suggesting insect cell infection.

Observation of the hijacking repertoire of *E. muscae* in houseflies showed that host manipulation lasts for ca. two hours, beginning within four hours of sunset six days post infection. Furthermore, we observed that few individual flies did not succumb to the infection and die on the sixth day after exposure, and that these flies displayed behaviours aberrant to uninfected flies (Supplementary Figure 1). A delayed death for few individuals could be an adaptive bet-hedging strategy by the pathogen(s) so that not all flies die at the same time in nature, which may increase the chance of transmission. On the contrary, the difference in time of death may also represent individual differences between flies in immune response or temporal variation in time of *E. muscae* infection leading to individual variation in disease progression between flies. The predictability of manipulated behaviour observations was central to specimen collection for collection of RNA samples that would allow us to identify candidate host and fungal genes involved in this extended phenotype (Naundrup *et al*., 2022). Mapping of RNA reads to insect and fungal genomes showed a decreased in read numbers mapping to the insect host with increasing fungal infection. This is in correlation with the logistic growth and build-up of *E. muscae* biomass inside house fly hosts during the course of infection (Hansen & De Fine Licht, 2017), and also observed for *E. muscae* infecting fruit flies (Elya *et al*., 2018). A decrease in reads mapping to the host genome is congruent to other fungus-insect systems, which suggested the parasites were proliferating in the insect heads during and after manipulation (Hughes *et al*., 2011; de Bekker *et al*., 2015; Will *et al*., 2020). However, we find that it is not the fungal, but a viral parasite which is proliferating in insect heads during infection (Figure 6a). This finding does not negate the importance of fungal SSPs and other candidate genes, which may be present in higher numbers outside fly heads since the fungus is predominantly found in the abdomen as it utilises host fatty tissue and freely available sugars (Brobyn & Wilding, 1983).

Analyses of insect and fungal differentially expressed genes (DEGs) in housefly heads identified putative candidate genes likely involved for inducing the summiting behaviour in *E. muscae*-infected houseflies. Specifically, there was an upregulation in genes enriched in immune functions (defence response to bacterium; innate immune response; regulation of Rho protein signal transduction) and response to stress (Supplementary Figure S1) in infected flies (IC), which suggests that the houseflies recognize the *E. muscae* infection. The fungus *E. muscae* proliferates as wall-less cells to evade the host immune system (Latgé *et al*., 1988; Boomsma *et al*., 2014), however the immune evasion appears not to be 100% as different components of the house fly immune response appear to be activated as also observed in *E. muscae* infected fruit flies (Elya *et al*., 2018). Enrichment in steroid hormone mediated signalling pathway, odorant binding, and nuclear steroid receptor activity among down-regulated genes during summiting, may suggest interference with the hosts ecdysone to mediate host immune responses (Zhu *et al*., 2021). During summiting, most enriched gene sets in IC were oppositely expressed (Supplementary Figure S1,2), which indicates a significant shift in host cellular homeostasis. Upregulation in transporters, steroid hormone mediated signalling pathway, cell junction, and postsynaptic membrane suggest that for summiting behaviour the host may be transporting fungal or viral products which enable the manipulated phenotype through increased ability for signal transmission. During PB, upregulation of genes enriched in sensory perception of smell and olfactory receptor activity may be involved in the extension of the proboscis through stimulation of the host’s olfactory system (Amrein & Thorne, 2005). By WG, downregulation in gene sets linked to physiological cellular metabolism likely indicates that the host houseflies were approaching death.

Fungal candidate genes which may be involved in summiting disease were identified through differential gene expression analysis. In particular, three genes were uniquely upregulated during summiting behaviour and downregulated again immediately after during proboscis extension, which indicates that these genes may play a key role in this manipulation phenotype. Of these three genes were two SSPs of unknown function, one containing a transmembrane protein, and the gene HBVY01013035, which encodes for a homologue of the *egt* summiting gene in behaviourally manipulating baculoviruses (Hoover *et al*., 2011; Han *et al*., 2015). Knockout of *egt* in the baculovirus *Lymantria dispar* multiple nucleopolyhedrovirus (LdMNPV) and *Spodoptera exigua* MNPV led to the *L. dispar* and *S. exigua* hosts dying on container bottoms as opposed to at elevated positions (Hoover *et al*., 2011). In insects, the enzyme encoded by *egt* inactivates the steroid hormone 20-hydroxyecdysone (20E) which regulates molting and metamorphosis in insects. However, the expression of 20E was not tested and we cannot infer on whether *egt* contributes to 20E suppression or to other mechanisms. A decrease in the expression of the enzyme encoded in *Sad* which catalyses 2.22-Dideoxy-edysone into ecdysone may suggest disruption of the molting hormone biosynthesis pathway, in particular the innate immune pathways in which ecdysone plays numerous roles (Haag *et al*., 1988). The EPF *Metarhizium rileyi* inactivates host ecdysone through the expression of ecdysteroid-22-oxidase, which play a role in mediating host responses to fungal infection through reducing host antimicrobial gene expression (Zhu *et al*., 2021). In *E. muscae*, the decrease in expression of *Sad* suggests a decreased catalysis and thus availability of ecdysone, thus reducing the host immune defence, which is observed in gene enrichment analyses of housefly genes during summiting behaviour.

Changes in the ecdysone receptor complex seen in this study (Figure 5b) may not be involved in summiting behaviour *per se*, as is seen in baculovirus-infected summiting in lepidopteran hosts via the secretion of *egt* to deactivate the steroid hormone 20E and delay molting (Hoover *et al*., 2011; Han *et al*., 2015). This pathway controls moulting and metamorphosis of insects (Niwa & Niwa, 2014), but as the houseflies used in this study are adults, the changes in the molting hormone pathway are unlikely to cause summiting behaviour, at least not through the same mechanisms. However, ecdysteroids in adult *Drosophila melanogaster* are involved in reproduction and sexual behaviours (e.g. egg and sperm production), circadian rhythm and neuronal functions (e.g. clock gene expression and neuronal activity and plasticity of clock neurons), and stress responses (e.g. reproductive physiology, sleep homeostasis and innate immunity) (Uryu *et al*., 2015; Yamanaka, 2021). This suggests that the disruption of genes involved in enzymatic activity of the insect molting hormones may be influencing the timing of *E. muscae* manipulations to occur before sunset and highlight the potential convergent evolution of mechanisms to induce summiting behaviours in insects between viral and fungal pathogens.

In all major fungal groups, there are viruses that can infect and replicate within fungal cells (Ghabrial *et al*., 2015). These mycoviruses are known to be present in insect, plant and human infecting fungal pathogens (Kotta-Loizou, 2021; Lerer & Shlezinger, 2022). Mycoviral infections exert a range of fitness costs to their pathogenic fungal host, from negative, to neutral and to positive effects, such as through modified virulence (reduced and enhanced), adaptations to new environments, mycotoxin production, and beneficial and detrimental effects (reduction or increase in virulence) (Kotta-Loizou, 2021; Lerer & Shlezinger, 2022). In houseflies infected with *E. muscae*, there has been found to be an almost ubiquitous symbiosis between the fungus and a +ssRNA iflavirus (Coyle et al. 2018; Myers *et al*., 2020; **Chapter 6)**. As iflaviruses are mainly known to be insect infecting, it is tempting to speculate that the fungus originally acquired the virus through horizontal transmission. One limiting factor for mycoviruses is usually that the lack of extracellular transmission to hosts due to the fungal cell wall that generally does not allow for the direct uptake or secretion of viruses, and due to the lack of proteins encoded for viral entry and spread indicative of intracellular life cycles, although exceptions do exist (Liu *et al*., 2016; Kotta-Loizou & Coutts, 2017a; Helenius, 2018). During the within-host life cycle, *E. muscae* proliferates as protoplasts, cells that do not have a cell wall. This likely affects the transmissibility of EVs into the host and may have facilitated the original fungal adoption of an insect virus. The viral permeability of the fungal protoplasts could allow the virus to systemically infect the insect through the use of fungal movement proteins. A similar mechanism is seen in one plant and one fungal RNA virus in which co-evolved mutualism has led to synergistic interactions between both viruses for horizontal transmission (Bian *et al*., 2020).

In other EPFs-insect systems, mycoviruses have been shown to alter fungal virulence (Kotta-Loizou & Coutts, 2017b), or alter volatile substances to attract fungus-eating flies (Liu *et al*., 2016). *E. muscae* uses volatile compounds to increase transmission during sporulation, through attracting necrophiliac male flies to attempt to copulate with the sporulating cadaver (Naundrup *et al*., 2022). As EVs appear to replicate in both host and fungal cells, it would be possible for EVs to influence volatile expression to increase transmission of both fungi and virus, as is seen in fungus-eating *Lycoriella ingenua* (Liu *et al*., 2016). Unfortunately, performing genetic manipulation in *E. muscae* and its sensitivity *in vitro* to antibiotics has hindered the removal of the virus to investigate EV function. Thus, the role that the virus plays in this tri-Kingdom interactions remains unknown.

The main question around EV function is whether it play a role in the observed manipulated behaviours. Baculoviruses are known to cause summiting disease and hyperactivity in their lepidopteran larvae hosts (Gasque *et al*., 2019; Wang & Hu, 2019). However, baculoviruses are dsDNA viruses and much more complex than EVs, which leads to how EVs could manipulate an insect host through the use of two polyproteins, although +ssRNA virus manipulation has been recorded (Dheilly *et al*., 2015). The parasitoid *Dinocampus coccinellae* injects an egg within its coccinellid host, *Coleomegilla maculate*. After *ca.* three weeks, the prepupa egresses and spins a cocoon between the legs of the host, during which time the host guards the parasitoid from predation. This bodyguard behaviour was attributed to the parasitoid’s doing, however Dheilly et al. (2015) found that an iflavirus, *Dinocampus coccinellae* paralysis virus (DcPV), which is transferred during oviposition replicates within the host nervous tissues after prepupa egression. Furthermore, after the adult wasp emerges, the ladybeetle’s normal behaviour returns, which also coincides with a decrease in viral titres. Leading to the conclusion that DcPV is likely responsible for the bodyguard behaviour, and not the parasitoid (Dheilly *et al*., 2015). An additional trait of an *E. muscae* infection that has attracted notice before is the formation of a clear liquid droplet at the tip of the proboscis of the zombie fly during proboscis manipulation (Elya *et al*., 2018; Elya & De Fine Licht, 2021). The purpose or cause of this excretion has not been studied, however is speculated to be a result of the increased pressure inside the fly caused by fungal growth. However, this phenotypic response has been reported in Chinese oak silkmoth (*Antheaea pernyi*) larvae infected with an iflavirus (ApIV) which cause *A. pernyi* vomiting disease (Geng *et al*., 2014). There is thus ample correlational indications for the importance of EVs in the manipulation of flies infected with *E. muscae,* but until causative evidence is available many of these patterns remain speculative.

## Conclusions

Psychomotor behaviours are governed by fine relationships between cognitive functions and physical activity, the severe incoordination of limbs during summiting suggests that the housefly cognitive functions are being affected, resulting in the observed psychomotor retardation (Iliadi *et al*., 2016). We identify multiple candidate fungal genes with involvement in summiting behaviour, but also highlight the need to further investigate the function of the symbiotic iflavirus found in *E. muscae*. Furthermore, we identify an *E. muscae egt* homologue significantly upregulated during summiting. *Egt* is known to be involved in summiting in baculoviruses, suggesting potential convergent evolution of mechanisms to induce summiting behaviours in insects between viral and fungal pathogens. Taken together, the results suggest that *E. muscae* is behaviourally manipulating its host by secreting small effector proteins and viral particles which interact with host proteins involved in neurological functions and disorders, and signal proteins involved in inflammation. Finally, observations in this study about the systemic presence of MdEV open up yet more questions as to whether the exhibited moribund phenotypes are truly an artefact of fungal origin.

## Methods

### 1. Fly stocks and fungus cultures

We used houseflies, *Musca domestica* (RNAseq using wild-type strain 772a, provided by University of Aarhus, Denmark; Virus quantifications using houseflies provided by MD-Terraristik, Germany) for experiments in this study. Pupated flies were sexed and sorted within 4 days of pupation and fed with ad libitum food (1:1 ratio caster sugar:semi skimmed milk powder) and demineralised water using a 10ml vial plugged with cotton. Fly culture and experiments were conducted at 20°C on a 12:12 light:dark cycle) in a non-humidified room. *Entomophthora muscae* isolates KVL21-40 and KVL22-01 were used to continuously infect healthy flies and were maintained at the same conditions in *in vivo* cultures (see Appendix 1 for more details on *E. muscae* infections).

### 2. Housefly infections

Three housefly cadavers newly killed (1-4 hours post death) by *E. muscae* were used to infect ten healthy male houseflies. The three cadavers, of which were a minimum of one male and one female, were fixed head first in 5% water agar in medicine cups. Ten adult male houseflies of 4-7 days old were placed in each medicine cups for 24 hours with 1 mg of food (as mention in section 1) in a saturated humid chamber. The cups were perforated for aeration and placed upside down to allow for the spore showers by the infected cadavers (Appendix 1). For the control treatment, 3 uninfected houseflies were euthanised by five minute exposure to -5°C. These dead uninfected houseflies replaced the sporulating cadavers used in the infected treatments. After 24 hours, the flies were anaesthetised with carbon dioxide before being transferred to larger pots with ad libitum food and water (under the same conditions as mentioned in section 1) until being sampled for RNA extraction five days later.

### 3. RNAseq sampling time points

Behaviour observations led to the selection of three manipulated behaviours of interest for RNA extractions. These known characteristic behaviours were a) summiting, b) proboscis extension and c) wing raising (Krasnoff *et al*., 1995; Elya & De Fine Licht, 2021). As the duration of these behaviours vary between individuals (pers. obs.; Krasnoff et al., 1995), individuals were sampled at points based on the subtleties of the manipulation process exhibited by the host houseflies during these behaviours of interest (see Supplementary information for a detailed overview of the sampling time points and the associated behaviours). Briefly, ‘summiting’ samples were collected when tremors, and locomotor retardation and incoordination were severely affected. For flies having summited, their proboscides pulsate (extending and retracting in larger movements over time) and bodies’ bob to touch mouthparts to the substrate surface (see Video S…). Samples were collected for ‘proboscis extension’ behaviour once proboscis pulsation stopped and the proboscides were in full extension, but before being fixed to the substrate. The ‘wing raise’ behaviour were defined when flies affixed by their proboscides had slowly unfolded their wings before raising them and were collected when wings were unfolded. In addition to these behaviourally manipulating samples, we collected samples from uninfected flies as negative control in parallel to the sampling of behaviourally manipulated flies. Furthermore, we collected infected flies to be used as infected control nine hours before subjected darkness (4-6 hours before summiting behaviour).

### 4. RNA extraction and sequencing

At these specific points, flies decapitated and immediately flash frozen in liquid nitrogen. For the uninfected treatment, the flies were placed in 5°C for ca. 2min prior to being decapitated and flash frozen. All specimens and RNA extractions were stored at -80°C.

RNA was extracted and purified using a Qiagen RNeasy plant mini kit (catalogue number 74904) from three pooled housefly heads with added lysis incubation at 56°C for 3 minutes, and phenol/chloroform and chloroform steps. Total RNA was then reversed transcribed to cDNA and used to prepare libraries that were sequenced using DNA Nanoball™ sequencing (DNBSeq™) technology. Library preparation and DNBSeq were performed by the Beijing Genomics Institute (BGI) Europe (Ole Maaloes Vej 3, 2200 Copenhagen N, Denmark).

### 5. Bioinformatic analysis

Raw read quality was checked with FastQC version 0.11.9 (https://www.bioinformatics.babraham.ac.uk/projects/fastqc/) and trimmed using trimmomatic version 0.38 (Bolger *et al*., 2014) with the following specifications: LEADING:5 TRAILING:5 SLIDINGWINDOW:4:15 MINLEN:40 -threads 20. The quality of the trimmed reads was assessed with FastQC. Following this quality and trimming steps, all RNAseq samples were processed as described below.

For housefly genes, we mapped the trimmed reads using STAR version 2.7.9a (Dobin *et al*., 2013) to the *Musca domestica* (aabys) genome (version 63) and associated structural annotation file downloaded from VectorBase on 18^th^ April 2023 (https://vectorbase.org/vectorbase/app/downloads/Current_Release/Mdomesticaaabys/). For the STAR index file, we specified the options --genomeSAindexNbases 13 and –sjdbOverhang 99. For the alignment, we used the options --outSAMtype BAM Unsorted, -- outReadsUnmapped Fastx, --chimSegmentMin 30, --chimOutType WithinBAM, -- twopassMode Basic and --peOverlapNbasesMin 5. Reads were left unsorted so that the files containing the unmapped reads could be analysed for the other organisms. For the reads that mapped to the housefly genome, samtools version 1.16 (Li *et al*., 2009) was used to sort the .bam files to be summarised for expressed gene counting using featureCounts version 2.0.2 (Liao *et al*., 2014). MultiQC was subsequently used for quality control of all of the steps prior to proceeding to differential gene expression analysis (Ewels *et al*., 2016).

For fungal genes, we mapped the reads that remained unmapped to the housefly genome to the *Entomophthora muscae* transcriptome (accession: HBVY01000001) using Kallisto version 0.46 (Bray *et al*., 2016). Again, samtools and MultiQC were run with the same options and following the same steps as for the housefly reads (Li *et al*., 2009; Ewels *et al*., 2016).

As we were aware of the presence of a mycoviral symbiont (Coyle *et al*., 2018; Myers *et al*., 2020), coined “Entomophthovirus” (EV), we used the genomes of eight EVs as a single reference for mapping (Supplementary Table S4). The genomes used for mapping were assembled using Trinity and originated from *E. muscae* infected houseflies (x1 genome), cabbage flies (*Delia radicum*) (x1 genome) and drosophilids (x6 genomes). Reads were pseudomapped to these viral reference genomes using Kallisto (Bray *et al*., 2016).

### 6. Differential gene expression analysis

The genes count tables created using featureCounts (Liao *et al*., 2014) and Kallisto (Bray *et al*., 2016) for host and fungal reads were normalised then analysed using DESeq2 (Love *et al*., 2014) in R (R Core Team, 2022). Due to the small temporal variation at which individuals were sampled (Figure 1), one summiting sample could not be clearly delineated from ‘infected control’ samples and was removed from subsequent analyses. Differentially expressed genes with a log-2 fold > 2 and with an adjusted p-value < 0.05 were extracted for downstream analyses.

### 7. Enrichment analysis of house fly genes

Analyses for gene enrichment during the behavioural time points was performed using the Gene Set Enrichment Analysis software (GSEA) using gene ontology (GO) terms as gene set on count data normalised in DESeq2. The enrichment tests were run for 1000 gene set permutations on all four pairwise comparisons for each behaviour phenotype and its previous time point, i.e. infected control Vs healthy, summiting Vs infected control, proboscis extension Vs summiting, and wing raising Vs proboscis extension. All gene set enrichments with a false discovery rate (FDR) cut-off of 5% were selected as recommended by GSEA for gene set permutation analysis. Gene set enrichments were visualised in Cytoscape using Enrichment Map.

### 8. Identification of an upregulated *E. muscae* UDP encoded gene

Differential expression analysis of *E. muscae* genes revealed that HBVY01013035, a 519 amino acid sequence which encodes for UDP-glucoronosyl and UDP-glucosyl transferase (UDP) and a transmembrane protein, is upregulated specifically during summiting behaviour. As the baculovirus ecdysteroid UDP-glucosyltransferase (*egt*) gene is known to be involved in summiting behaviour in some systems, we performed a BLASTp search Altschul et al., 1997) of all non-redundant *baculoviridae* CDS translations on GenBank to identify *egt* homology of gene HBVY01013035. The protein sequences from the top 10 blastp hits based on the lowest e-value, one sequence of a known MNPV *egt* involved in summiting, five selected fungal *egt* sequences (*Colletotrichum* spp.), and a hypothetical UDP from *Massospora cicadina* were downloaded from NCBI. The protein sequences were aligned using MAFFT v7 (Katoh & Standley, 2013) before being trimmed using BMGE (Criscuolo & Gribaldo, 2010). The LG+G model was selected for phylogenetic analysis was selected using SMS v1.8.1 (Lefort *et al*., 2017) and used in maximum likelihood analysis using PhyML (Guindon *et al*., 2010) on the NGPhylogeny.fr online workflow (Lemoine *et al*., 2019). The phylogenetic tree was visualised using the programme FigTree version 1.4.4 (Rambaut, 2018).

The sequences of the *E. muscae* UDP, the 11 baculovirus *egt*, and the *M. cicadina* hypothetical UDP were aligned in MAFFT (Katoh & Standley, 2013). The conserved domain architectures of the *E. muscae* UDP protein sequence were searched for against the conserved domain database (Wang *et al*., 2023) using CD-Search (Marchler-Bauer & Bryant, 2004), and the identified active site residues for the glycosyltransferase domain were extracted from the aligned sequences. Protein sequence motifs for active site residues were obtained using MEME v5.5.4 (Bailey *et al*., 2006).

### 9. Isolation and identification of a new *E. muscae* isolate and its viruses

Due to a collapse of the isolate KVL21-40 in the lab, a new isolate was collected from houseflies caught in a cow byre in Osted, Denmark (55.549204; 11.948141), kept under rearing conditions in the lab and isolated in liquid culture from sporulating flies in GLEN media (Beauvais and Latgé, 1988; **Chapter 8**). The new *E. muscae* isolate was labelled KVL22-01.

Fungal DNA was extracted from actively growing cultures in liquid GLEN medium. 2 mL of liquid culture was centrifuged at 1400 rpm for 10 minutes. The supernatant was removed and the fungal pellet was washed once with 1 mL MilliQ H_2_O, resuspended and centrifuged at 1400 rpm for 10 minutes, then at 4000 rpm for 5 minutes. 500 μL of MilliQ H_2_O was removed and then samples were subjected to beads beating in a TissueLyser (Qiagen) at 30 beats per second for 2 minutes. Fungal DNA was extracted with a Qiagen DNEasy Plant mini kit (catalogue number 69204) according to the manufacturer’s instructions. The internal transcribed spacer (ITS) region was amplified with a polymerase chain reaction (PCR), using primers ITS1 5’-TCCGTAGGTGAACCTGCGG and ITS4 5’-TCCTCCGCTTATTGATATGC (White *et al*., 1990). Thermocycling was performed as follows: 95℃ for 3 minutes, then 40 cycles of 95℃ for 30 seconds, 55 ℃ for 30 seconds, 72℃ for 60 seconds, followed by 72℃ for 7 minutes. PCR products were assessed on a 0.8% agarose-TBE buffer gel and purified using an illustra™ GFX PCR DNA and Gel Band Purification Kit from Cytiva following manufacturer’s protocol. The purified PCR products were sequenced using Sanger sequencing (Macrogen EUROPE).

For the virus, the RNA-dependent RNA polymerase (RdRp) regions of the viruses associated with the KVL21-40 (formerly KVL21-01) and KVL21-41 (infected *Drosophila* from Denmark) *E. muscae* isolates were obtained using the ORFfinder (blastORF) and blastp against the UniProtKB/Swiss-Prot(swissprot) database. The resulting RdRp sequences were checked and confirmed using palmid (version 0.0.5), resulting in a 920 nucleotide DNA or 120-122 amino acid protein sequence. Using the RdRp region of the viral genomes obtained from *E. muscae* infected houseflies and Danish drosophilids (**Chapter 6**), virus RdRp regions were aligned using Clustal Omega (Sievers & Higgins, 2018) to select sequence regions of interest to design PCR primers using Primer3 (Koressaar & Remm, 2007; Untergasser *et al*., 2012). The following primer sets were developed from the infected housefly (isolate KVL21-40): MdEV_F 5’ GGATACATTGAAGGATGAGCGG and MdEV_R 5’ CATGAGCCTCTAGTAAAGCGC for a product size of 904 nucleotides. And the following set was designed from the infected *Drosophila* (isolate KVL21-42) virus RdRp: DspEV_F 5’ AAGGATGAACGCAAAAGCCA and DspEV_R 5’ CGCTTTTCCACACCCATTGA for a product size of 838 nucleotides.

Subsequently, total RNA was extracted from uninfected flies, wing raising flies infected with KVL21-40, isolate KVL22-01 from liquid media and distilled water as negative control. RNA was purified then reverse transcribed to cDNA following manufacturers’ protocols using Deoxyribonuclease I, Amplification Grade kit (catalogue number 18068-015) and iScript cDNA Synthesis Kit (catalogue number 1708891), respectively. The cDNA samples were amplified by PCR using the following protocol: : 94℃ for 3 minutes, then 28 cycles of 94℃ for 60 seconds, 52 (DspEv) to 54 (MdEV) ℃ for 60 seconds, 72℃ for 90 seconds, followed by 72℃ for 5 minutes. The PCR products were assessed on a 2% agarose-TBE buffer gel and purified using an illustra™ GFX PCR DNA and Gel Band Purification Kit from Cytiva following manufacturer’s protocol. The purified KVL22-01 PCR products were sequenced using Sanger sequencing (Macrogen EUROPE).

### 10. Analysis of virus sequences

The ITS and RdRp sequences from the KVL22-01 isolate were cleaned and aligned in Geneious version 2023.2.1. Forward and reverse ITS sequences were aligned and a consensus sequence was generated to confirm this isolate to be *E. muscae* using the Basic Local Alignment Search Tool (BLAST) (Altschul et al., 1997).

For the virus, forward and reverse sequences were aligned within primer sets (MdEV and DspEV) to make a consensus sequence. The MdEv and DspEV consensus sequences were aligned using default Geneious Prime version 2023.2.1 multiple alignment, revealing that both sequences had a pairwise identity of 82.9% and so were treated as two different viruses in the following steps. The RdRp region of the two new viruses were determined using National Centre for Biotechnology Information (NCBI) ORFfinder (https://www.ncbi.nlm.nih.gov/orffinder/) and BLASTp. The resulting RdRp sequence was checked and confirmed using palmid version 0.0.5 (Edgar *et al*., 2021; Babaian & Edgar, 2022), resulting in a 920 nucleotide DNA or 122 amino acid (aa) protein sequence.

We used the genome of the virus from the KVL21-40 to predict the structure of the virus since we did only had the RdRp regions of the MdEV1 and MdEV2 viruses isolated from *E. muscae* isolate KVL22-01. Genome structure was predicted using InterProScan to analyse and identify viral proteins (Paysan-Lafosse *et al*., 2023).

### 11. Virus phylogeny

The RNA-dependent RNA polymerases (RdRps) of iflaviruses and related viruses were used for phylogenetic analysis. Accession numbers of representative iflaviruses from officially recognised species were obtained from the International Committee on Taxonomy of Viruses (ICTV) Report chapter on *Iflaviridae* (https://ictv.global/report/chapter/iflaviridae/iflaviridae) (Valles *et al*., 2017).

Accession numbers of iflaviruses not yet officially classified were obtained using the BLASTp with the newly sequenced mycoviruses as queries. Accession numbers of representative secoviruses from officially recognised species were obtained from the ICTV Report chapter on *Secoviridae* (https://ictv.global/report/chapter/secoviridae/secoviridae) (Fuchs *et al*., 2022).

All amino acid sequences of RdRPs were obtained from the NCBI Protein Database. The Molecular Evolutionary Genetics Analysis (MEGA) 11 software (Tamura et al., 2021) was used for phylogenetic analysis. A multiple alignment of RdRP amino acid sequences was produced using the MUltiple Sequence Comparison by Log-Expectation (MUSCLE) algorithm (Edgar 2004) as implemented by MEGA. The Le & Gascuel (LG) model with frequencies (+F) and gamma distributed (G) rates among sites was determined by MEGA to be the most appropriate for the RdRp amino acid sequence alignment. All positions in the multiple alignment with less than 50% site coverage were eliminated. A maximum likelihood (ML) phylogenetic tree was created by MEGA using the bootstrap method (100 replicates) as a phylogeny test.

### 12. Quantification of viral load in different host body parts

Total RNA was extracted from homogenised head, thorax and abdomen of flies collected during ‘wing raising’ behavioural manipulation using a Qiagen RNeasy plant mini kit (catalogue number 74904) according to the manufacturer’s instructions with an added incubation step at 56°C for 10 minutes in lysis buffer. RNA was purified using the Deoxyribonuclease I, Amplification Grade kit (catalogue number 18068-015; ThermoFisher Scientific) following the manufacturer’s protocol. Purified RNA was then reverse-transcribed into 200 ng complementary DNA (cDNA) using the iScript cDNA Synthesis Kit (catalogue number 1708891; BIO-RAD) and following the manufacturer’s protocol.

Primers for real-time polymerase chain reaction (RT-PCR) were designed based on the RNA-dependent RNA polymerase (RdRp) region of the two identified KVL22-01 EVs using primer3 (Koressaar & Remm, 2007; Untergasser *et al*., 2012). Primers for *Musca domestica* Entomophthovirus 1 (MdEV1) were MdEV1_605_F 5’-CCGAATTTCTTAAGCGAGGGT and MdEV1_704_R 5’-ACCCATTGTGTGATGTCCTCA, for a product size of 100 nt. For *Musca domestica* Entomophthovirus 2 (MdEV2), the primer set was MdEV2_53_F 5’-CCGAATGGGCTGGATTAGTG and MdEV2_148_R 5’-CAGCATTAAATCCTGGTCCGA, for a product size of 96 nt. The RdRp region was amplified with a PCR using the MdEV1 and MdEV2 primer sets and using a thermal gradient from 52 to 64°C. Thermocycling was performed as follows: 95℃ for 3 minutes, then 40 cycles of 95℃ for 30 seconds, 52-64℃ for 30 seconds, 72℃ for 60 seconds, followed by 72℃ for 7 minutes. PCR products were assessed on a 2% agarose-TAE buffer gel and purified using an illustra™ GFX PCR DNA and Gel Band Purification Kit (GELifeSciences). Samples with the strongest gel electrophoresis band (59°C for MdEV1 and 54°C for MdEV2) were selected and purified PCR products’ DNA concentrations were measured using QUBIT DNA Broad Range Kit. These two selected samples were used to produce standard curves for absolute quantification of viral cDNA copy numbers down the line for the samples of interest through cycle threshold value comparisons (Supplementary Figure S6-S9).

For samples, flies were infected as previously mentioned and sampled 0 (uninfected), 24 (1 dpi), 96 (4 dpi) and 135 (6 am dpi) hours after infection. Flies were sampled also during proboscis extension to represent a behaviour manipulation time point, 15 hours after darkness initiated for a sample of ca. 12 hours into sporulation, and from an actively growing fungal culture 11 days after transfer to new media. For the insect samples, the head, thorax and abdomens were separated before being homogenised. For the fungal culture, 2 mL of liquid culture was centrifuged at 4000 rpm for 5 minutes. The supernatant was removed and the fungal pellet was washed once with 500 μL MilliQ H_2_O, resuspended and centrifuged at 4000 rpm for 5 minutes to remove media. The pellet was resuspended in 500 μL MilliQ H_2_O and subject to beads beating in a TissueLyser (Qiagen) at 30 beats per second for a further 5 minutes. Total RNA extraction, purification and reverse-transcription were performed on all samples as mentioned above for samples used to generate standard curves. Some samples had too low and RNA concentration to be reverse transcribed into 200 ng of cDNA, these were reverse transcribed into 100 ng instead. Thermocycling was performed as mentioned above using 1 μL of cDNA per sample to quantify the viral load using the specific MdEV1 and MdEV2 qPCR primers (Supplementary Figure S6,S8). Triplicates of each sample were used and the standard curve efficiency was considered to remain unchanged during qPCR. Thereafter, the molecular weight of the samples were calculated using the corresponding standard curves (for MdEV1 or MdEV2) to quantify the concentration of target cDNA. From these, copy numbers were calculated using the following equation:

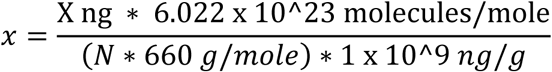

Where: X = amount of amplicon (ng); N = length of dsDNA amplicon; 660 g/mol = average mass of 1 bp dsDNA; 6.022 x 1023 = Avogadro’s constant; and 1 x 10^9^ = Conversion factor (https://eu.idtdna.com/pages/education/decoded/article/calculations-converting-from-nanograms-to-copy-number).

Samples that were reverse transcriber into 100 ng of cDNA were assumed to half the copy numbers of 200 ng samples. Thus, the calculated viral copy numbers of the 100 ng samples were multiplied by two. In total, three flies were used for time 0 (uninfected control) and sporulation, and six to eight were used for each of the other time points. For 1 dpi heads, only four samples had over 10 ng/μL.

### 13. Data Analyses of viral load

All analyses were performed in R version 4.2.2 (R Core Team, 2022). A Shapiro–Wilk test of normality was used to test for normal distribution of the log-transformed data. Since the data were not normally distributed, generalized linear models (GLM) were built using the lme4 package. The GLMs were compared based on the values of the Akaike information criterion (AIC), and the models with the lowest AIC were selected. An Analysis of Variance ANOVA was used to estimate the significant effects of the models.

Firstly, virus ID was used as fixed factor in a model, which showed a significant difference in mean viral titre of both viruses, thus the viruses were analysed independently. For each virus, the models including an interaction between body part and dpi were best fitting the data and so used a fixed effects. Due to the significance of body part, dpi and their interaction, the variables were joined together for pairwise comparisons using post-hoc analyses using the tukey method with the Bonferroni correction to account for multiple testing in the ‘emmeans’ package.

### 14. Protein-Protein interactions

To understand how host-pathogen proteins may be interacting, we used D-SCRIPT (v 0.2.2) to generate predicted interactions between host-fungus, host-virus, and fungus-virus. The *M. domestica* (aabys) proteome (version 63 downloaded from VectorBase) was used for host proteins interactions. The nucleotide sequences of the KVL21-40 virus was translated to protein sequences using transdecoder (Haas, BJ. https://github.com/TransDecoder/TransDecoder), revealing 114 open reading frames (ORFs) longer than 30 amino acids. Predicted ORFs were identified using InterProScan (Paysan-Lafosse *et al*., 2023), revealing two Iflavirus polyproteins, including a 40 amino acid long protein with a transmembrane domain located within the protease encoded by the 3’-proximal ORF.

For the fungus, DGE analysis revealed 17 fungal genes that were identified as candidates involved in summiting. Of these, 7 contained a signal peptide in their functional annotation, which were translated into 7 protein sequences using transdecoder (Haas, BJ. https://github.com/TransDecoder/TransDecoder). To verify if these proteins with SignalP domains were small secreted proteins (SSPs), effector predictions were made using EffectorP version 3.0 (Sperschneider & Dodds, 2022). Moreover, EffectorP predicts whether proteins are cytoplasmic or apoplastic effectors, and these localisation types were verified using ApoplastP version 1.0 (Sperschneider *et al*., 2018).

Using the seven candidate *E. muscae* SSPs, the three virus ORFs and the *M. domestica* (aabys) proteome, we tested every interspecific combination of virus, parasite and host protein to generate predicted interactions with D-SCRIPT (v 0.2.2) (n = 131,789 combinations tested for fungus-host; n = 21 for virus-fungus; n = 56,481 for virus-host) (Sledzieski *et al*., 2021). For computational time, we limited the input *M. domestica* proteins to only include those ≤ 2,000 amino acids in length (removing 723 of 19,550 sequences). We used D-SCRIPT in prediction mode with the default human-protein pretrained model and settings. GO term enrichment of fly genes predicted to be involved in PPI with virus or fungal proteins were analysed using ShinyGO version 0.77 with an FDR cut-off of 0.05 (Ge *et al*., 2020).

## Notes

### Competing Interest Statement

The authors have declared no competing interest.

## References

Altschul, S. F., Madden, T. L., Schäffer, A. A., Zhang, J., Zhang, Z., Miller, W. & Lipman D. J. 1997. Gapped BLAST and PSI-BLAST: a new generation of protein database search programs. Nucleic Acids Research, 25: 3389–3402.

Amrein, H. & Thorne, N. 2005. Gustatory Perception and Behavior in *Drosophila melanogaster*. Current Biology 15: R673–R684.

Babaian A. & Edgar R.C. 2022. Ribovirus classification by a polymerase barcode sequence. PeerJ 10:e14055.

Bailey, T.L., Williams, N., Misleh, C. & Li, W.W. 2006. MEME: discovering and analyzing DNA and protein sequence motifs. Nucleic Acids Research 34: W369–W373.

Bauer, A., Haine, E.R., Perrot-Minnot, M.-J. & Rigaud, T. 2005. The acanthocephalan parasite *Polymorphus minutus* alters the geotactic and clinging behaviours of two sympatric amphipod hosts: the native *Gammarus pulex* and the invasive *Gammarus roeseli*. Journal of Zoology 267: 39–43. Cambridge University Press.

Beckerson, W.C., Krider, C., Mohammad, U.A. & de Bekker, C. 2023. 28 minutes later: investigating the role of aflatrem-like compounds in *Ophiocordyceps* parasite manipulation of zombie ants. Animal Behaviour 203: 225–240.

Bian, R., Andika, I.B., Pang, T., Lian, Z., Wei, S., Niu, E., et al. 2020. Facilitative and synergistic interactions between fungal and plant viruses. Proceedings of the National Academy of Sciences 117: 3779–3788.

Bolger, A.M., Lohse, M. & Usadel, B. 2014. Trimmomatic: a flexible trimmer for Illumina sequence data. Bioinformatics 30: 2114–2120.

Boomsma, J.J., Jensen, A.B., Meyling, N.V. & Eilenberg, J. 2014. Evolutionary Interaction Networks of Insect Pathogenic Fungi. Annual Review of Entomology 59: 467–485.

Bray, N.L., Pimentel, H., Melsted, P. & Pachter, L. 2016. Near-optimal probabilistic RNA-seq quantification. Nat Biotechnol 34: 525–527.

Brobyn, P.J. & Wilding, N. 1983. Invasive and developmental processes of *Entomophthora muscae* infecting houseflies (Musca domestica). Transactions of the British Mycological Society 80: 1–8.

Brown, D.W., Butchko, R.A.E., Busman, M. & Proctor, R.H. 2012. Identification of gene clusters associated with fusaric acid, fusarin, and perithecial pigment production in *Fusarium verticillioides*. Fungal Genetics and Biology 49: 521–532.

Chase, W.R., Zhaxybayeva, O., Rocha, J., Cosgrove, D.J. & Shapiro, L.R. 2020. Global cellulose biomass, horizontal gene transfers and domain fusions drive microbial expansin evolution. New Phytologist 226: 921–938.

Coyle, M.C., Elya, C.N., Bronski, M. & Eisen, M.B. 2018. Entomophthovirus: An insect-derived iflavirus that infects a behavior manipulating fungal pathogen of dipterans. bioRxiv.

Criscuolo, A. & Gribaldo, S. 2010. BMGE (Block Mapping and Gathering with Entropy): a new software for selection of phylogenetic informative regions from multiple sequence alignments. BMC Evolutionary Biology 10: 210.

Dawkins, R. 1982. The extended phenotype: The long reach of the gene. Oxford University Press.

De Bekker, C., Beckerson, W.C. & Elya, C. 2021. Mechanisms behind the Madness: How Do Zombie-Making Fungal Entomopathogens Affect Host Behavior To Increase Transmission? mBio 12: 10.1128/mbio.01872-21.

de Bekker, C., Ohm, R.A., Loreto, R.G., Sebastian, A., Albert, I., Merrow, M., et al. 2015. Gene expression during zombie ant biting behavior reflects the complexity underlying fungal parasitic behavioral manipulation. BMC Genomics 16: 620.

De Fine Licht, H.H., Edwards, S. & Elya, C. 2023. Evolutionary ecology of an obligate and behaviorally manipulating insect-pathogenic fungus, Entomophthora muscae. Authorea.

De Fine Licht, H.H., Jensen, A.B. & Eilenberg, J. 2017. Comparative transcriptomics reveal host-specific nucleotide variation in entomophthoralean fungi. Molecular Ecology 26: 2092–2110.

Dheilly, N.M., Maure, F., Ravallec, M., Galinier, R., Doyon, J., Duval, D., et al. 2015. Who is the puppet master? Replication of a parasitic wasp-associated virus correlates with host behaviour manipulation. Proceedings of the Royal Society B: Biological Sciences 282: 20142773.

Dobin, A., Davis, C.A., Schlesinger, F., Drenkow, J., Zaleski, C., Jha, S., et al. 2013. STAR: ultrafast universal RNA-seq aligner. Bioinformatics 29: 15–21.

Döhner, K., Nagel, C.-H. & Sodeik, B. 2005. Viral stop-and-go along microtubules: taking a ride with dynein and kinesins. Trends in Microbiology 13: 320–327.

Edgar, R. C. 2004. MUSCLE: multiple sequence alignment with high accuracy and high throughput. Nucleic Acids Research, 32, 1792–1797.

Edgar, R.C., Taylor, J., Lin, V., et al. 2022. Petabase-scale sequence alignment catalyses viral discovery. Nature 602: 142–147.

Elya, C. & De Fine Licht, H.H. 2021. The genus *Entomophthora*: bringing the insect destroyers into the twenty-first century. IMA Fungus 12: 34.

Elya, C., Lavrentovich, D., Lee, E., Pasadyn, C., Duval, J., Basak, M., et al. 2023. Neural mechanisms of parasite-induced summiting behavior in ‘zombie’ *Drosophila*. eLife 12: e85410.

Elya, C., Lok, T.C., Spencer, Q.E., McCausland, H., Martinez, C.C. & Eisen, M. 2018. Robust manipulation of the behavior of *Drosophila melanogaster* by a fungal pathogen in the laboratory. eLife 7: e34414.

Evans, H.C. 1989. 9 - Mycopathogens of Insects of Epigeal and Aerial Habitats. In: Insect-fungus Interactions (N. Wilding, N. M. Collins, P. M. Hammond, & J. F. Webber, eds), pp. 205–238. Academic Press, London.

Ewels, P., Magnusson, M., Lundin, S. & Käller, M. 2016. MultiQC: summarize analysis results for multiple tools and samples in a single report. Bioinformatics 32: 3047–3048.

Fuchs, M., Hily, J.-M., Petrzik, K., Sanfaçon, H., Thompson, J.R., van der Vlugt, R., et al. 2022. ICTV Virus Taxonomy Profile: *Secoviridae*. Journal of General Virology 103: 001807.

Gasque, S.N., van Oers, M.M. & Ros, V.I. 2019. Where the baculoviruses lead, the caterpillars follow: baculovirus-induced alterations in caterpillar behaviour. Current Opinion in Insect Science 33: 30–36.

Ge, S.X., Jung, D. & Yao, R. 2020. ShinyGO: a graphical gene-set enrichment tool for animals and plants. Bioinformatics 36: 2628–2629.

Geng, P., Li, W., Lin, L., Miranda, J.R. de, Emrich, S., An, L., et al. 2014. Genetic Characterization of a Novel Iflavirus Associated with Vomiting Disease in the Chinese Oak Silkmoth *Antheraea pernyi*. PLoS ONE 9: e92107.

Ghabrial, S.A., Castón, J.R., Jiang, D., Nibert, M.L. & Suzuki, N. 2015. 50-plus years of fungal viruses. Virology 479–480: 356–368.

Gofton, A.W., Blasdell, K.R., Taylor, C., Banks, P.B., Michie, M., Roy-Dufresne, E., et al. 2022. Metatranscriptomic profiling reveals diverse tick-borne bacteria, protozoans and viruses in ticks and wildlife from Australia. Transboundary and Emerging Diseases 69: e2389–e2407.

Guindon, S., Dufayard, J.-F., Lefort, V., Anisimova, M., Hordijk, W. & Gascuel, O. 2010. New Algorithms and Methods to Estimate Maximum-Likelihood Phylogenies: Assessing the Performance of PhyML 3.0. Systematic Biology 59: 307–321.

Guittard, E., Blais, C., Maria, A., Parvy, J.-P., Pasricha, S., Lumb, C., et al. 2011. CYP18A1, a key enzyme of *Drosophila* steroid hormone inactivation, is essential for metamorphosis. Developmental Biology 349: 35–45.

Haag, T., Hetru, C., Kappler, C., Marie Moustier, A., A. Hoffmann, J. & Luu, B. 1988. Study on the biosynthesis of ecdysone: part IV(1):Synthesis of high specific activity (3H2∼22,23)∼2,22-dideoxyecdysone tissue distribution of the C-22 hydroxylase in *Locusta migratoria*. Tetrahedron 44: 1397–1408.

Han, Y., Van Houte, S., Drees, G.F., Van Oers, M.M. & Ros, V.I.D. 2015. Parasitic manipulation of host behaviour: baculovirus SeMNPV EGT facilitates tree-top disease in *Spodoptera exigua* larvae by extending the time to death. Insects 6: 716–731.

Hansen, A.N. & De Fine Licht, H.H. 2017. Logistic growth of the host-specific obligate insect pathogenic fungus *Entomophthora muscae* in house flies (*Musca domestica*). Journal of Applied Entomology 141: 583–586.

Helenius, A. 2018. Virus Entry: Looking Back and Moving Forward. Journal of Molecular Biology 430: 1853–1862.

Herbison, R.E.H. 2017. Lessons in Mind Control: Trends in Research on the Molecular Mechanisms behind Parasite-Host Behavioral Manipulation. Frontiers in Ecology and Evolution 5.

Hoover, K., Grove, M., Gardner, M., Hughes, D.P., McNeil, J. & Slavicek, J. 2011. A Gene for an Extended Phenotype. Science 333: 1401–1401.

Hughes, D.P., Andersen, S.B., Hywel-Jones, N.L., Himaman, W., Billen, J. & Boomsma, J.J. 2011. Behavioral mechanisms and morphological symptoms of zombie ants dying from fungal infection. BMC Ecology 11: 13.

Iliadi, K.G., Gluscencova, O.B. & Boulianne, G.L. 2016. Psychomotor Behavior: A Practical Approach in *Drosophila*. Frontiers in Psychiatry 7: 153.

Katoh, K. & Standley, D.M. 2013. MAFFT Multiple Sequence Alignment Software Version 7: Improvements in Performance and Usability. Molecular Biology and Evolution 30: 772–780.

Katsuma, S., Koyano, Y., Kang, W., Kokusho, R., Kamita, S.G. & Shimada, T. 2012. The Baculovirus uses a captured host phosphatase to induce enhanced locomotory activity in host caterpillars. PLOS Pathogens 8: e1002644.

Kikawada, T., Saito, A., Kanamori, Y., Nakahara, Y., Iwata, K., Tanaka, D., et al. 2007. Trehalose transporter 1, a facilitated and high-capacity trehalose transporter, allows exogenous trehalose uptake into cells. Proceedings of the National Academy of Sciences 104: 11585–11590.

Kokusho, R. & Katsuma, S. 2021. *Bombyx mori* nucleopolyhedrovirus *ptp* and *egt* genes are dispensable for triggering enhanced locomotory activity and climbing behavior in *Bombyx mandarina* larvae. Journal of Invertebrate Pathology 183: 107604.

Koressaar, T. & Remm, M. 2007. Enhancements and modifications of primer design program Primer3. Bioinformatics 23: 1289–1291.

Kotta-Loizou, I. 2021. Mycoviruses and their role in fungal pathogenesis. Current Opinion in Microbiology 63: 10–18.

Kotta-Loizou, I. & Coutts, R.H.A. 2017a. Mycoviruses in *Aspergilli*: A Comprehensive Review. Frontiers in Microbiology 8.

Kotta-Loizou, I. & Coutts, R.H.A. 2017b. Studies on the virome of the entomopathogenic fungus *Beauveria bassiana* reveal novel dsRNA elements and mild hypervirulence. PLOS Pathogens 13: e1006183.

Krasnoff, S.B., Watson, D.W., Gibson, D.M. & Kwan, E.C. 1995. Behavioral effects of the entomopathogenic fungus, *Entomophthora muscae* on its host *Musca domestica*: Postural changes in dying hosts and gated pattern of mortality. Journal of Insect Physiology 41: 895–903.

Lafferty, K.D. & Shaw, J.C. 2013. Comparing mechanisms of host manipulation across host and parasite taxa. Journal of Experimental Biology 216: 56–66.

Latgé, J.P., Eilenberg, J., Beauvais, A. & Prevost, M.C. 1988. Morphology of *Entomophthora muscae* protoplasts grown in vitro. Protoplasma 146: 166–173.

Lefort, V., Longueville, J.-E. & Gascuel, O. 2017. SMS: Smart Model Selection in PhyML. Molecular Biology and Evolution 34: 2422–2424.

Lemoine, F., Correia, D., Lefort, V., Doppelt-Azeroual, O., Mareuil, F., Cohen-Boulakia, S., et al. 2019. NGPhylogeny.fr: new generation phylogenetic services for non-specialists. Nucleic Acids Research 47: W260–W265.

Lerer, V. & Shlezinger, N. 2022. Inseparable companions: Fungal viruses as regulators of fungal fitness and host adaptation. Frontiers in Cellular and Infection Microbiology 12.

Li, H., Handsaker, B., Wysoker, A., Fennell, T., Ruan, J., Homer, N., et al. 2009. The Sequence Alignment/Map format and SAMtools. Bioinformatics 25: 2078–2079.

Liang, M., Du, S., Dong, W., Fu, J., Li, Z., Qiao, Y., et al. 2019. iTRAQ-based quantitative proteomic analysis of mycelium in different predation periods in nematode trapping fungus *Duddingtonia flagrans*. Biological Control 134: 63–71.

Liao, Y., Smyth, G.K. & Shi, W. 2014. featureCounts: an efficient general purpose program for assigning sequence reads to genomic features. Bioinformatics 30: 923–930.

Liu, S., Xie, J., Cheng, J., Li, B., Chen, T., Fu, Y., et al. 2016. Fungal DNA virus infects a mycophagous insect and utilizes it as a transmission vector. Proceedings of the National Academy of Sciences 113: 12803–12808.

Liu, X., Tian, Z., Cai, L., Shen, Z., Michaud, J.P., Zhu, L., et al. 2022. Baculoviruses hijack the visual perception of their caterpillar hosts to induce climbing behaviour thus promoting virus dispersal. Molecular Ecology 31: 2752–2765.

Love, M.I., Huber, W. & Anders, S. 2014. Moderated estimation of fold change and dispersion for RNA-seq data with DESeq2. Genome Biology 15: 550.

Lovett, B., Macias, A., Stajich, J.E., Cooley, J., Eilenberg, J., Licht, H.H. de F., et al. 2020. Behavioral betrayal: How select fungal parasites enlist living insects to do their bidding. PLOS Pathogens 16: e1008598.

Marchler-Bauer, A. & Bryant, S.H. 2004. CD-Search: protein domain annotations on the fly. Nucleic Acids Research 32: W327–W331.

Martin, S.W. & Konopka, J.B. 2004. Lipid Raft Polarization contributes to hyphal growth in *Candida albicans*. Eukaryotic Cell 3: 675–684.

Maure, F., Brodeur, J., Ponlet, N., Doyon, J., Firlej, A., Elguero, É., et al. 2011. The cost of a bodyguard. Biology Letters 7: 843–846.

Myers, J.M., Bonds, A.E., Clemons, R.A., Thapa, N.A., Simmons, D.R., Carter-House, D., et al. 2020. Survey of Early-Diverging Lineages of Fungi Reveals Abundant and Diverse Mycoviruses. mBio 11: 10.1128/mbio.02027-20.

Narváez-Barragán, D.A., Tovar-Herrera, O.E., Segovia, L., Serrano, M. & Martinez-Anaya, C. 2020. Expansin-related proteins: biology, microbe–plant interactions and associated plant-defense responses. Microbiology 166: 1007–1018.

Naundrup, A., Bohman, B., Kwadha, C.A., Jensen, A.B., Becher, P.G. & De Fine Licht, H.H. 2022. Pathogenic fungus uses volatiles to entice male flies into fatal matings with infected female cadavers. ISME J, 16, 2388–2397.

Niwa, R. & Niwa, Y.S. 2014. Enzymes for ecdysteroid biosynthesis: their biological functions in insects and beyond. Bioscience, Biotechnology, and Biochemistry 78: 1283–1292.

Obayashi, N., Iwatani, Y., Sakura, M., Tamotsu, S., Chiu, M.-C. & Sato, T. 2021. Enhanced polarotaxis can explain water-entry behaviour of mantids infected with nematomorph parasites. Current Biology 31: R777–R778.

O’Reilly, D.R., Brown, M.R. & Miller, L.K. 1992. Alteration of ecdysteroid metabolism due to baculovirus infection of the fall armyworm *Spodoptera frugiperda*: Host ecdysteroids are conjugated with galactose. Insect Biochemistry and Molecular Biology 22: 313–320.

Paysan-Lafosse, T., Blum, M., Chuguransky, S., Grego, T., Pinto, B.L., Salazar, G.A., et al. 2023. InterPro in 2022. Nucleic Acids Research 51: D418–D427.

Ponton, F., Otálora-Luna, F., Lefèvre, T., Guerin, P.M., Lebarbenchon, C., Duneau, D., et al. 2011. Water-seeking behavior in worm-infected crickets and reversibility of parasitic manipulation. Behavioral Ecology 22: 392–400.

Poulin, R. & Maure, F. 2015. Host manipulation by parasites: A look back before moving forward. Trends in Parasitology 31: 563–570.

Rambaut, A. (2018). FigTree-v1.4.4. http://treebioedacuk/software/figtree/ (accessed April 5, 2023).

R Core Team. (2022). R: A language and environment for statistical computing. R Foundation for Statistical Computing, Vienna, Austria. See https://www.R-project.org/.

Ros, V.I.D., van Houte, S., Hemerik, L. & van Oers, M.M. 2015. Baculovirus-induced tree-top disease: how extended is the role of egt as a gene for the extended phenotype? Molecular Ecology 24: 249–258.

Sievers, F. & Higgins, D.G. (2018) Clustal Omega for making accurate alignments of many protein sequences. Protein Science 27:135–145.

Shang, Y., Feng, P. & Wang, C. 2015. Fungi that infect insects: Altering host behavior and beyond. PLOS Pathogens 11: e1005037.

Singh, A. & Del Poeta, M. 2011. Lipid signalling in pathogenic fungi. Cellular Microbiology 13: 177–185.

Sledzieski, S., Singh, R., Cowen, L. & Berger, B. 2021. D-SCRIPT translates genome to phenome with sequence-based, structure-aware, genome-scale predictions of protein-protein interactions. Cell Systems 12: 969–982.e6.

Sperschneider, J. & Dodds, P.N. 2022. EffectorP 3.0: Prediction of apoplastic and cytoplasmic effectors in Fungi and Oomycetes. MPMI 35: 146–156.

Sperschneider, J., Dodds, P.N., Singh, K.B. & Taylor, J.M. 2018. ApoplastP: prediction of effectors and plant proteins in the apoplast using machine learning. New Phytologist 217: 1764–1778.

Tamura, K., Stecher, G. & Kumar, S. 2021. MEGA11: Molecular evolutionary genetics analysis version 11. Molecular Biology and Evolution 38: 3022–3027.

Takasuka, K., Yasui, T., Ishigami, T., Nakata, K., Matsumoto, R., Ikeda, K., et al. 2015. Host manipulation by an ichneumonid spider ectoparasitoid that takes advantage of preprogrammed web-building behaviour for its cocoon protection. Journal of Experimental Biology 218: 2326–2332.

Tixier, V., Bataillé, L., Etard, C., Jagla, T., Weger, M., DaPonte, J.P., et al. 2013. Glycolysis supports embryonic muscle growth by promoting myoblast fusion. Proceedings of the National Academy of Sciences 110: 18982–18987.

Untergasser, A., Cutcutache, I., Koressaar, T., Ye, J., Faircloth, B.C., Remm, M., et al. 2012. Primer3—new capabilities and interfaces. Nucleic Acids Research 40: e115.

Uryu, O., Ameku, T. & Niwa, R. 2015. Recent progress in understanding the role of ecdysteroids in adult insects: Germline development and circadian clock in the fruit fly *Drosophila melanogaster*. Zoological Letters 1: 32.

Valles, S.M., Chen, Y., Firth, A.E., Guérin, D.M.A., Hashimoto, Y., Herrero, S., et al. 2017. ICTV Virus Taxonomy Profile: *Iflaviridae*. Journal of General Virology 98: 527–528.

van Houte, S., Ros, V.I.D., Mastenbroek, T.G., Vendrig, N.J., Hoover, K., Spitzen, J., et al. 2012. Protein tyrosine phosphatase-induced hyperactivity is a conserved strategy of a subset of baculoviruses to manipulate lepidopteran host behavior. PLOS ONE 7: e46933.

van Houte, S., Ros, V.I.D. & van Oers, M.M. 2013. Walking with insects: molecular mechanisms behind parasitic manipulation of host behaviour. Molecular Ecology 22: 3458–3475.

Wang, J., Chitsaz, F., Derbyshire, M.K., Gonzales, N.R., Gwadz, M., Lu, S., et al. 2023. The conserved domain database in 2023. Nucleic Acids Research 51: D384–D388.

Wang, M. & Hu, Z. 2019. Cross-talking between baculoviruses and host insects towards a successful infection. Philosophical Transactions of the Royal Society B: Biological Sciences 374: 20180324.

White, T.J., Bruns, T., Lee, S. & Taylor, J. 1990. 38 – Amplification and direct sequencing of fungal ribosomal RNA genes for phylogenetics. In: PCR Protocols (M. A. Innis, D. H. Gelfand, J. J. Sninsky, & T. J. White, eds), pp. 315–322 Academic Press, San Diego.

Will, I., Beckerson, W.C. & de Bekker, C. 2023. Using machine learning to predict protein– protein interactions between a zombie ant fungus and its carpenter ant host. Scientific Reports 13: 13821.

Will, I., Das, B., Trinh, T., Brachmann, A., Ohm, R.A. & de Bekker, C. 2020. Genetic underpinnings of host manipulation by *Ophiocordyceps* as revealed by comparative transcriptomics. G3 (Bethesda) 10: 2275–2296.

Yamanaka, N. 2021. Chapter One - Ecdysteroid signalling in insects—From biosynthesis to gene expression regulation. In: Advances in Insect Physiology (M. E. Adams, ed), pp. 1–36. Academic Press.

Zhang, Y., Sampathkumar, A., Kerber, S.M.-L., Swart, C., Hille, C., Seerangan, K., et al. 2020. A moonlighting role for enzymes of glycolysis in the co-localization of mitochondria and chloroplasts. Nat Commun 11: 4509.

Zhu, S., Feng, X., Keyhani, N.O., Liu, Y., Jin, D., Tong, S., et al. 2021. Manipulation of host ecdysteroid hormone levels facilitates infection by the fungal insect pathogen, *Metarhizium rileyi*. Environmental Microbiology 23: 5087–5101.

