## Supplementary material for "Pathogenic fungus expresses effector proteins in combination with a symbiotic virus to behaviourally manipulate housefly hosts": suppl info

### Supplementary Methods

### Manipulated behaviours

#### 1. Summiting

The behaviour termed as ‘summiting’ is generally regarded as the act of when the infected host ascends to an elevated position, when it will inevitably die, and prior to performing the sequence of behaviours to come. This particular behaviour in the *E. muscae* infected houseflies is distinguished by a disruption in limb coordination during locomotion and the desire to climb. The climbing desire is noticeable as if the fly falls (from loss of locomotor skills), it will then attempt anew until reaching a desired height. If the fly does fall during ‘summiting’, the summiting behaviour will be visible for an extended duration compared to the immobile state it could be in if it is in the desired position early on in the ‘summiting’ manipulation. This means that the behaviour of summiting can be ongoing even if the characteristic act is not performed. It must also be noted that if the individual falls soon before the transition to the following behaviour then it generally remains immobile on the floor and proceeds to the next step (i.e. proboscis extension). Thus, samples for the behaviour category of ‘summiting’ were collected during the point at which locomotor skills were most severely disrupted and flies were still attempting to climb (after being knocked down or falling).

#### 2. Proboscis extension

Once the host fly has reached the desired elevation, it then undergoes the process of extending and affixing its proboscis to the substrate surface, termed ‘proboscis extension and affixation’. Prior to this manipulation execution, the host does not simply extend its proboscis once it has reached this height. If the fly does not fall and need to re-climb, the fly will remain in the desired death location immobile for an extended period until whatever molecular signals have been changed to initiate the proboscis to commence extending. The categorical manipulated behaviour ‘proboscis extension’ is (visible) initiated by the pulsation of the proboscis. This pulsation changes in magnitude but the proboscis retraction becomes lesser with the ongoing pulsation, until it remains fully extended (sometimes missing the net and poking through a whole (photo)). Once extended and in contact with the surface of the substrate, the proboscis begins to affix, sometimes with a clear liquid being produced out the mouth, and presumed to be held in place with a ‘fungal holdfast’. It is interesting to note that once the proboscis is affixed, the host fly is sometimes seen to appear to be pulling away. The samples collected for the 'proboscis extension' category were collected once the proboscis had stopped pulsating and was fully extended.

#### 3. Wing raising

Once the proboscis is affixed and the fly stops fighting the attachment, the fly again goes into a state of apparent immobility. Eventually the wings unfold from the resting position in one lengthy movement. Once the wings are unfolded, they begin to raise and the fly's body position prostrates as if in kowtow. This is the final visible behaviour manipulation, following which the fly remains immobile, dies then sporulates. The specific timing of death is not visible, but the fly main react to rough stimulus (grabbed by forceps) during the wing raising process.

The fly is sampled for the 'wing raising' behaviour at the point where the wing is unfolded but prior to the wings raising, again this behaviour occurs over different timeframes depending on the individual. It has not been determined whether the wing unfolding and elevation is due to molecular signals or fungal cells causing actively causing the muscles in the thorax related to flight to contract, as seen in the mandibles of ants infected with *Ophiocordyceps* (de Bekker et al., 2018). The wing raising would be explained by contraction of the indirect vertical muscles used to raise the wings during flight. As for the unfolding of the wings, this could be the same but in the anterior and/or posterior muscles. This could also provide an explanation for the kowtow seen following the wings being raised fully. Muscle contractions limited to thorax/wings, are there gene that code for contractions of muscles and how do they go from proboscis to wing unfolding to wing raising and then to kowtow?

#### 4. Pre-manipulation (infected control)

In order to have a molecular control to eliminate the gene expression of the fungal development, samples were also collected from individuals that were in a late stage of infection but within 12 hours of 'dawn'. This time was selected to ensure that the flies were fully infected and known to have *E. muscae* be undergoing changes from protoplasts to hyphal growth morphology (Elya and De Fine Licht, 2021; Annette Bruun-Jensen Thesis). These flies were collected simultaneously to the collection of the manipulated flies after five minute exposure to 5˚C for anaesthesia.

### Supplementary Figures

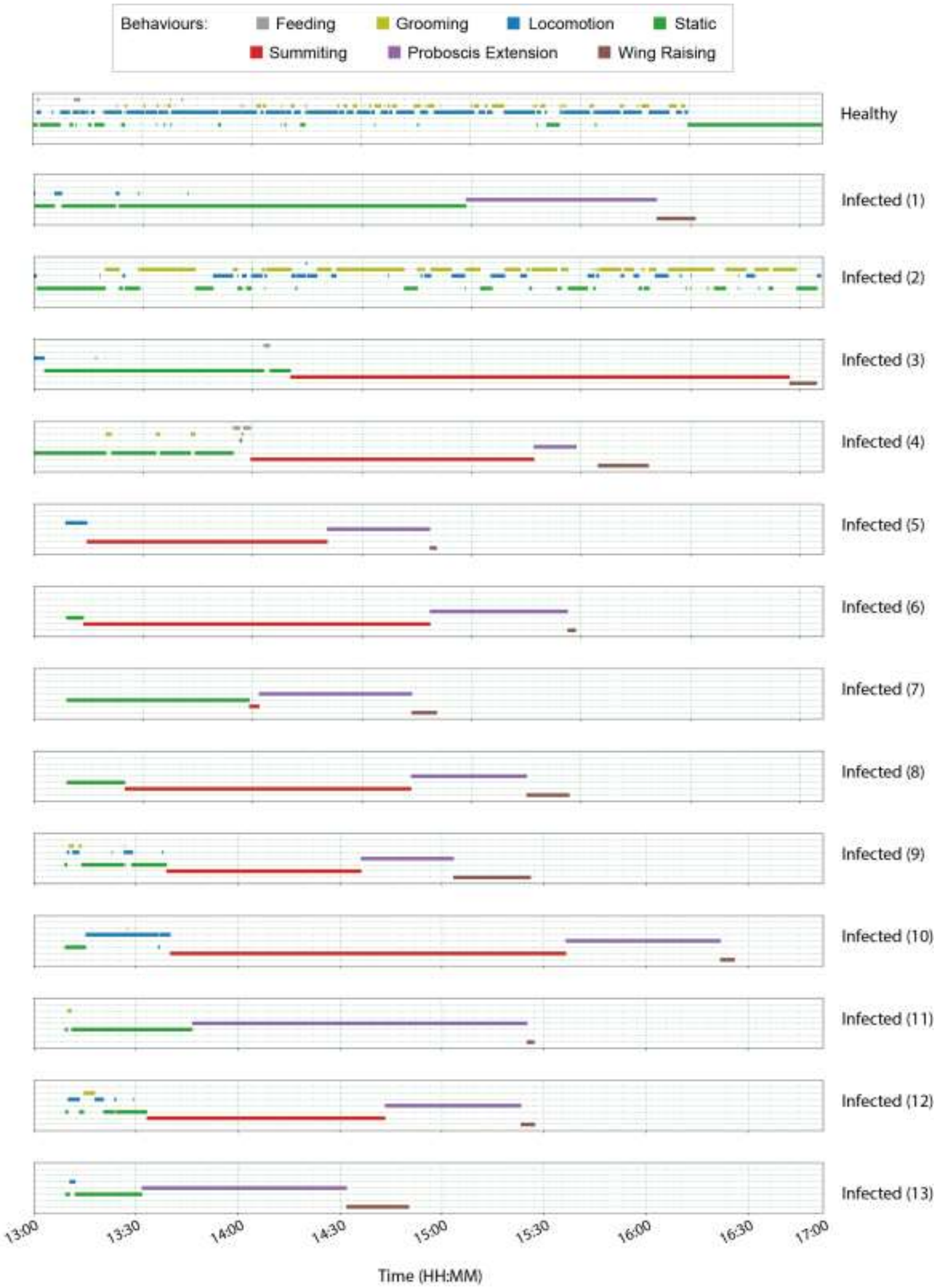


**Supplementary Figure S1.** Activity of houseflies infected and non-infected six day post infection.

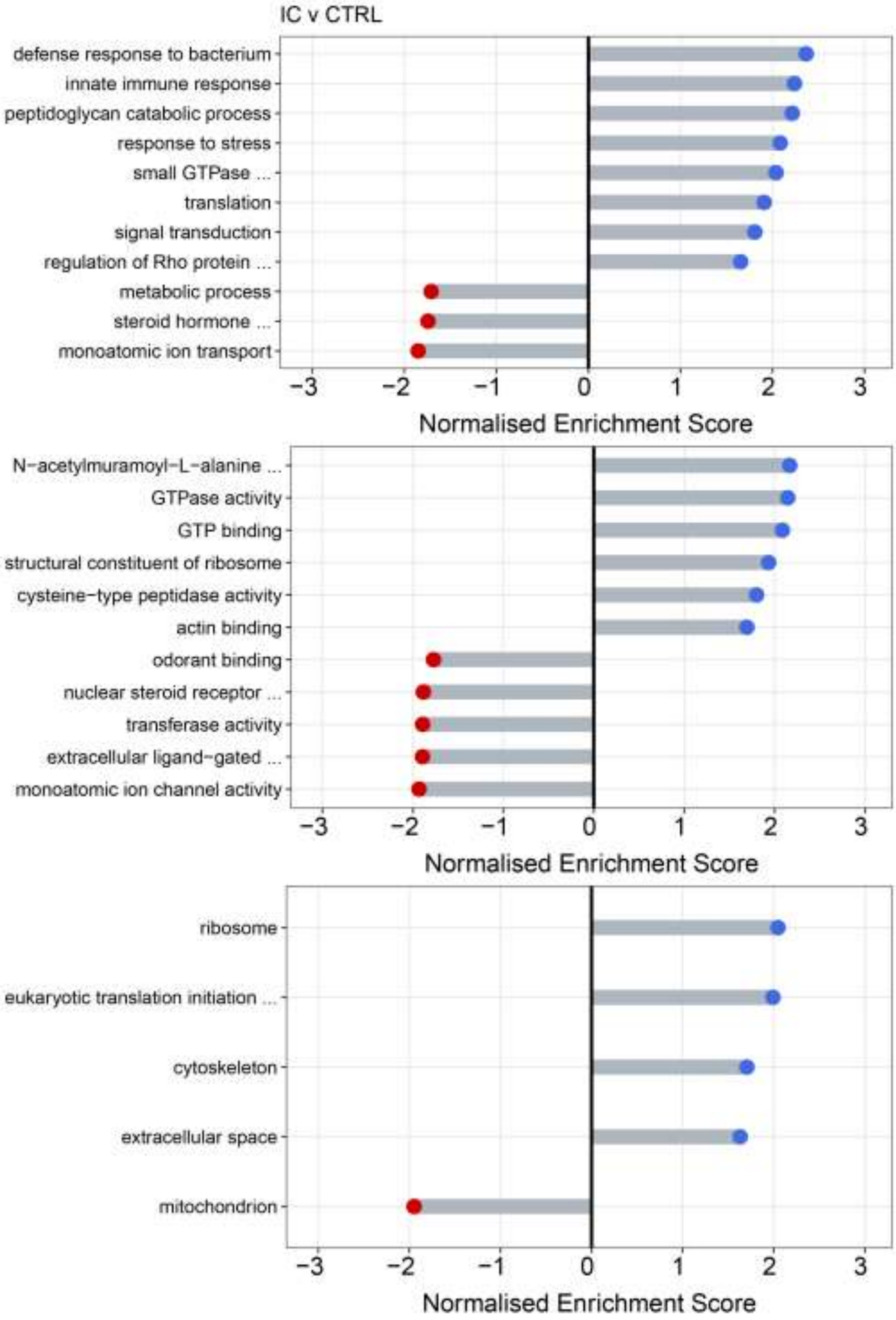


**Supplementary Figure S1.** Housefly host Gene Ontology term enrichment for infected control compared to uninfected flies. Top, middle and bottom graph represent enrichments for the biological processes, molecular function, and cellular components, respectively.

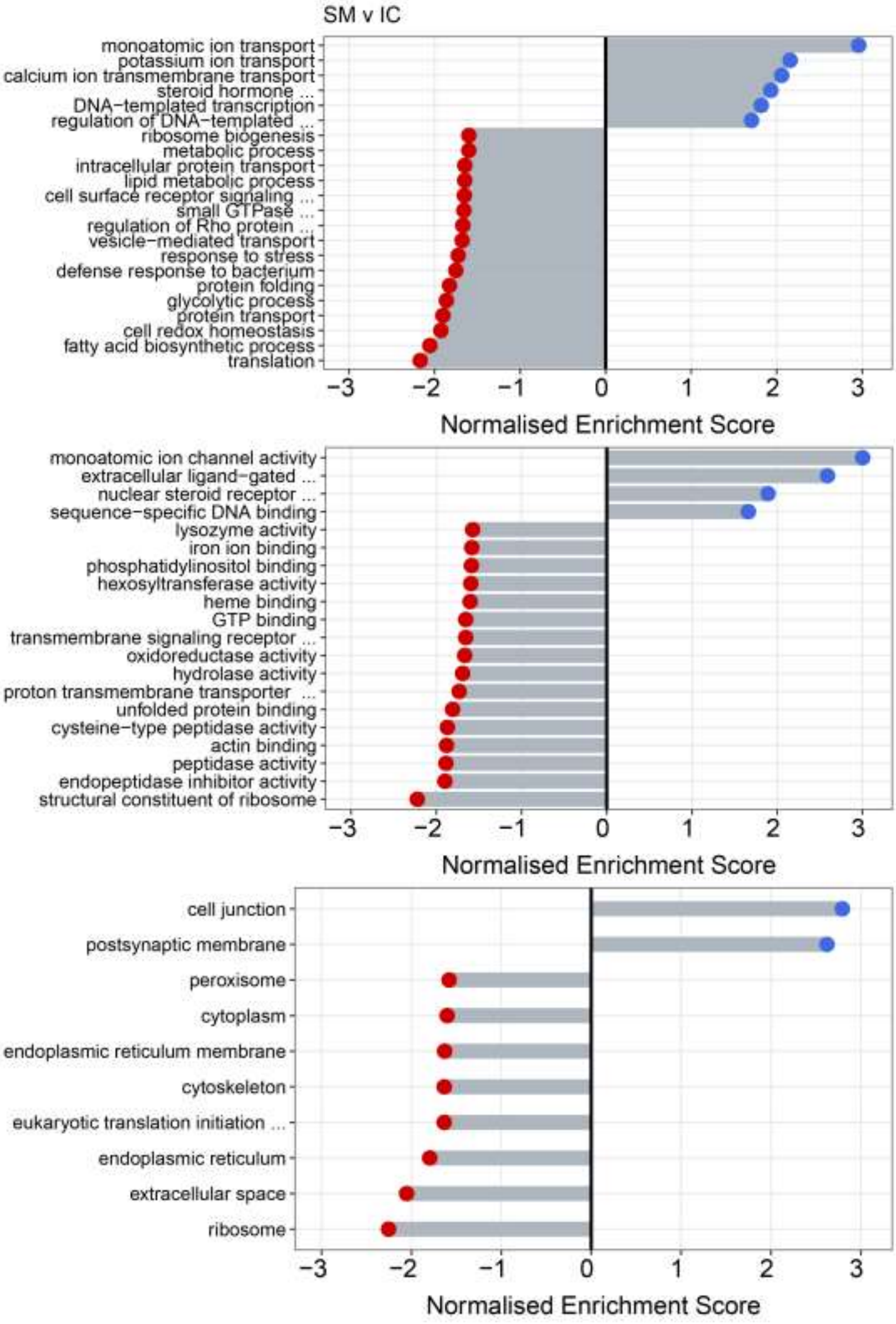


**Supplementary Figure S2.** Housefly host Gene Ontology term enrichment for summiting compared to infected control flies. Top, middle and bottom graph represent enrichments for the biological processes, molecular function, and cellular components, respectively.

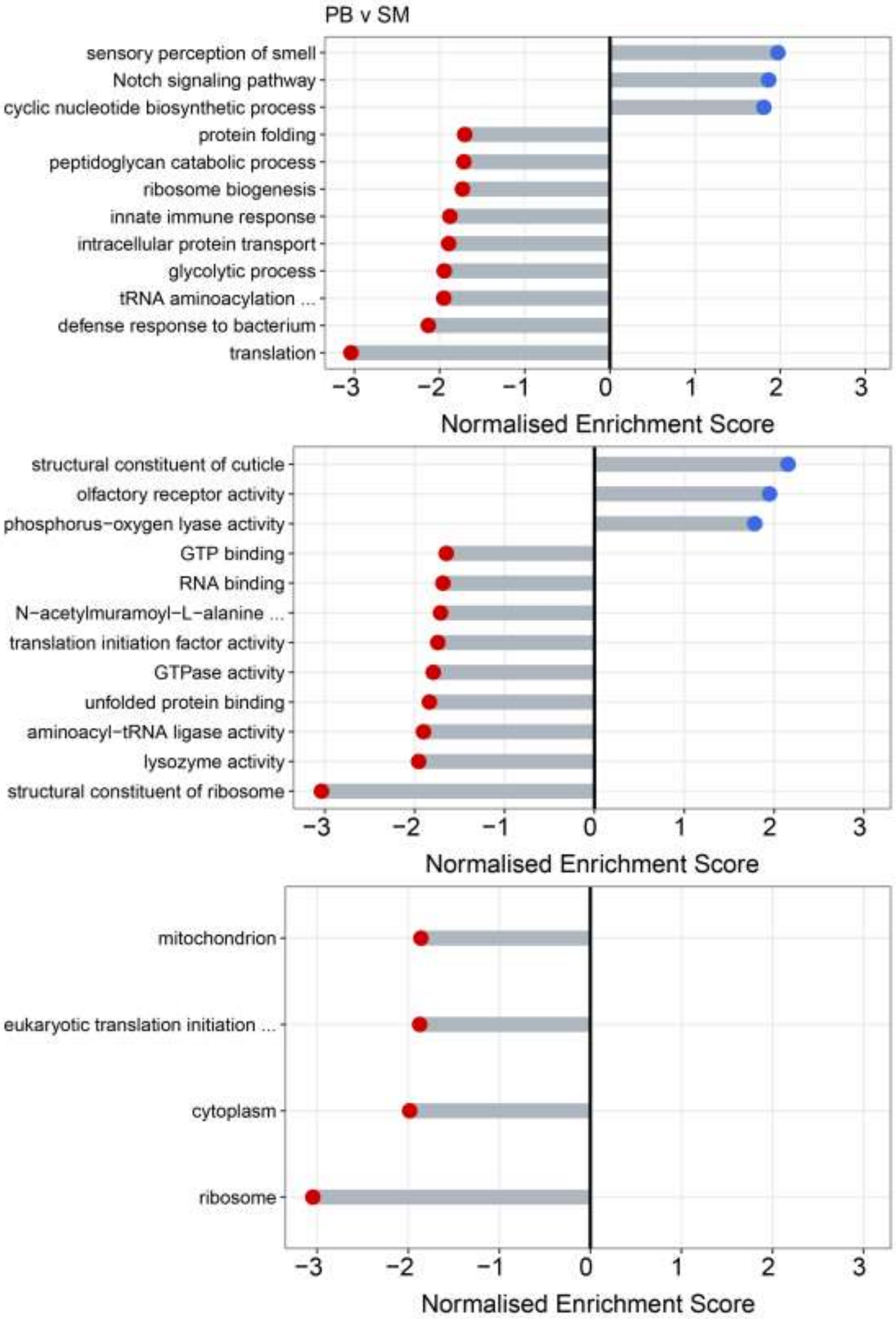


**Supplementary Figure S3.** Housefly host Gene Ontology term enrichment for proboscis extension compared to summiting flies. Top, middle and bottom graph represent enrichments for the biological processes, molecular function, and cellular components, respectively.

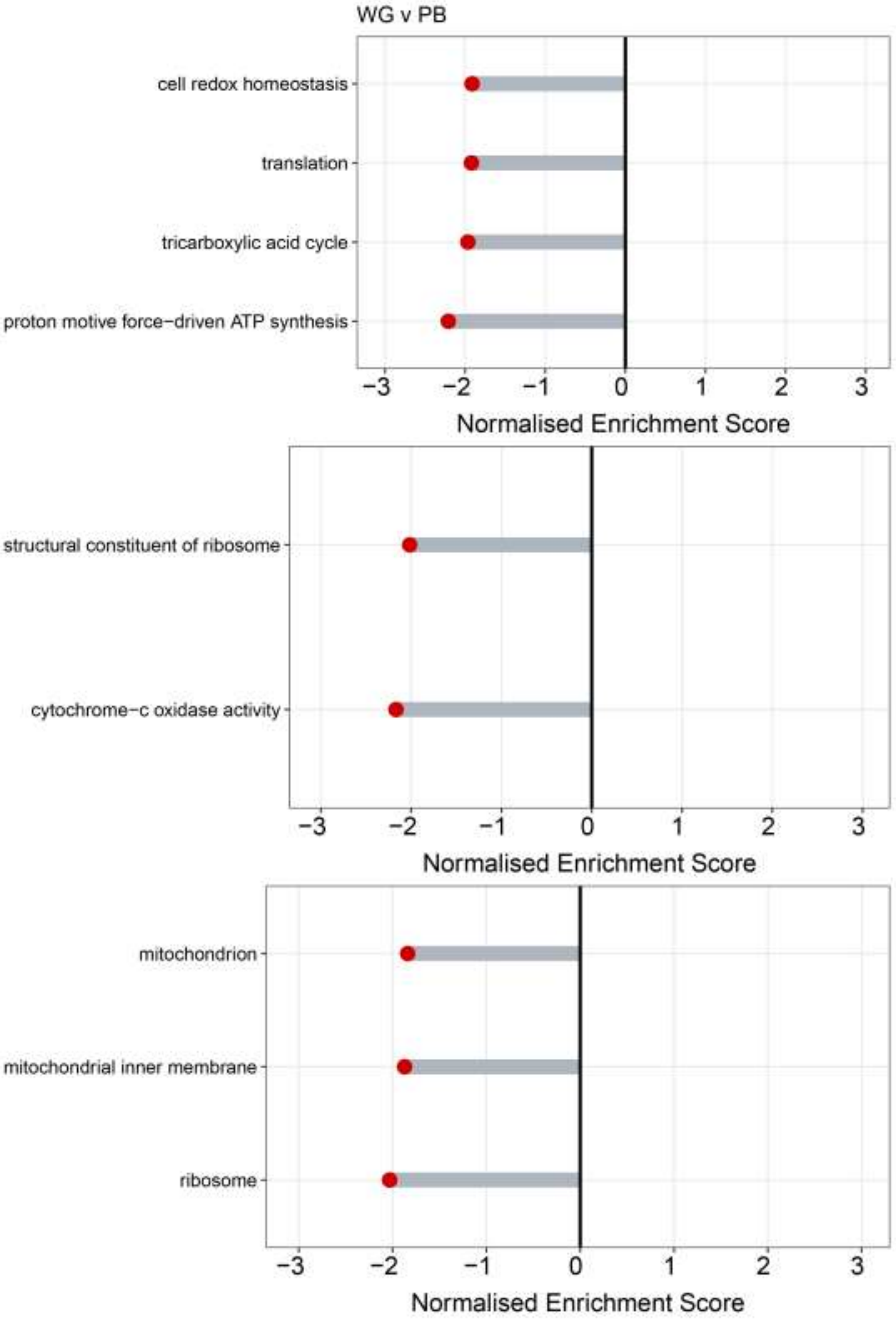


**Supplementary Figure S4.** Housefly host Gene Ontology term enrichment for wing raising compared to proboscis extension flies. Top, middle and bottom graph represent enrichments for the biological processes, molecular function, and cellular components, respectively.

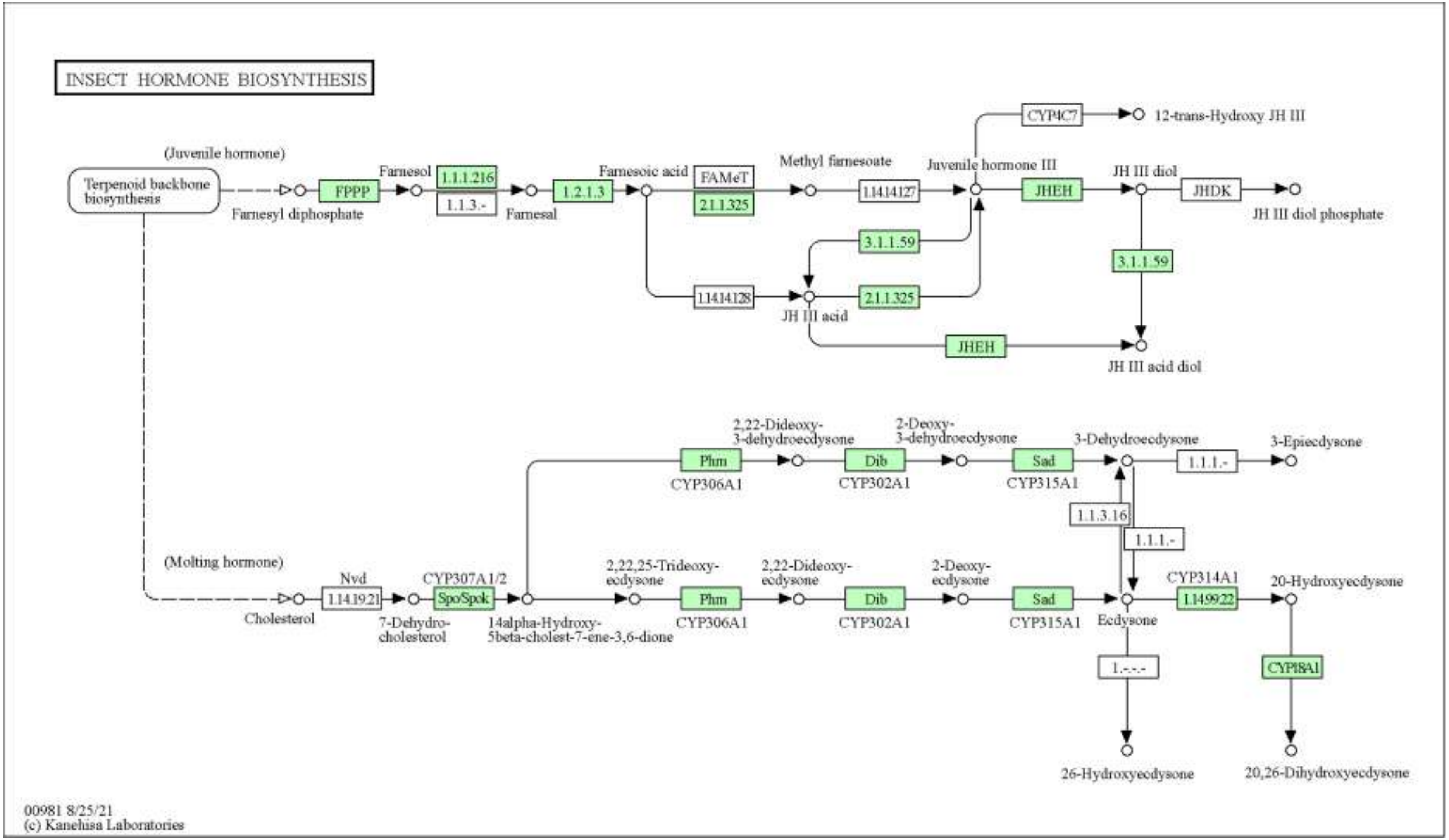


**Supplementary Figure S5.** Housefly insect hormone biosynthesis pathway map downloaded from the KEGG PATHWAY database (https://www.kegg.jp/kegg/pathway.html).

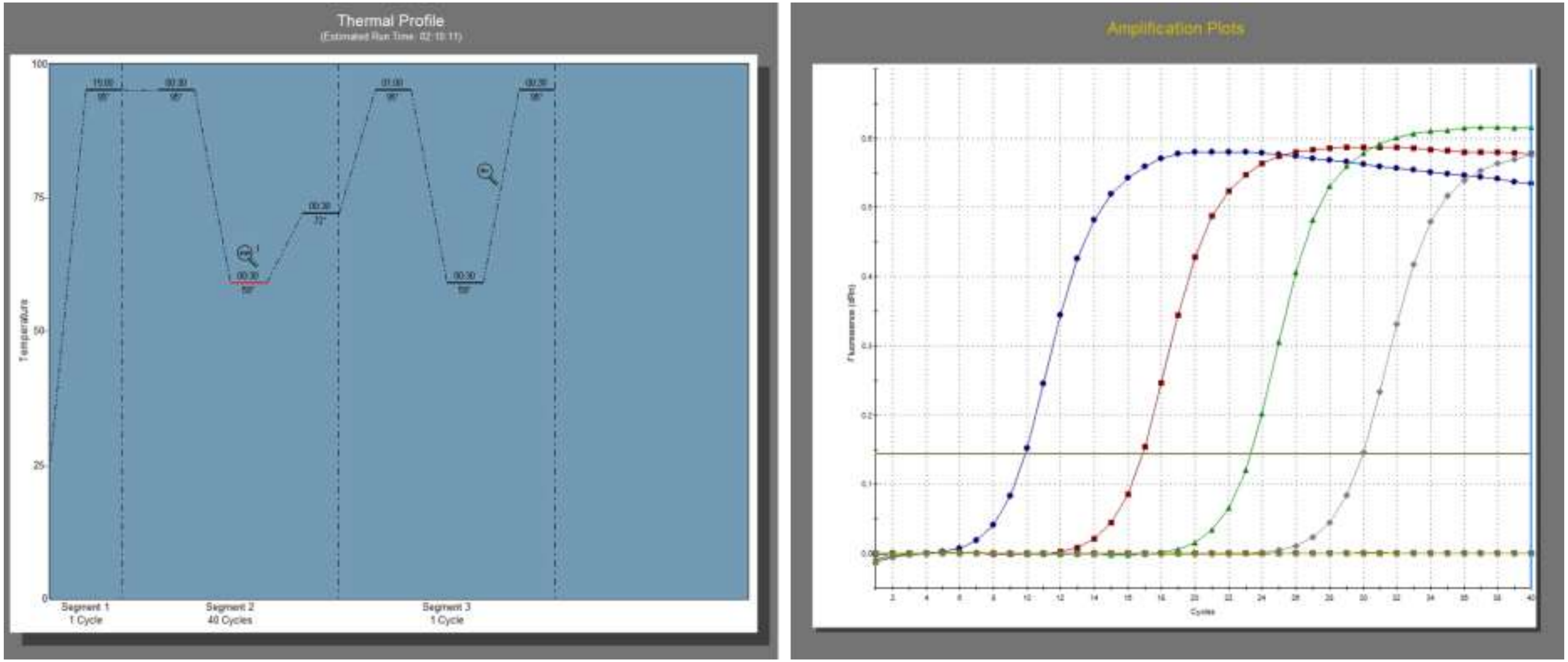


**Supplementary Figure S6.** Thermal profile and amplification plots for MdEV1 qPCRs.

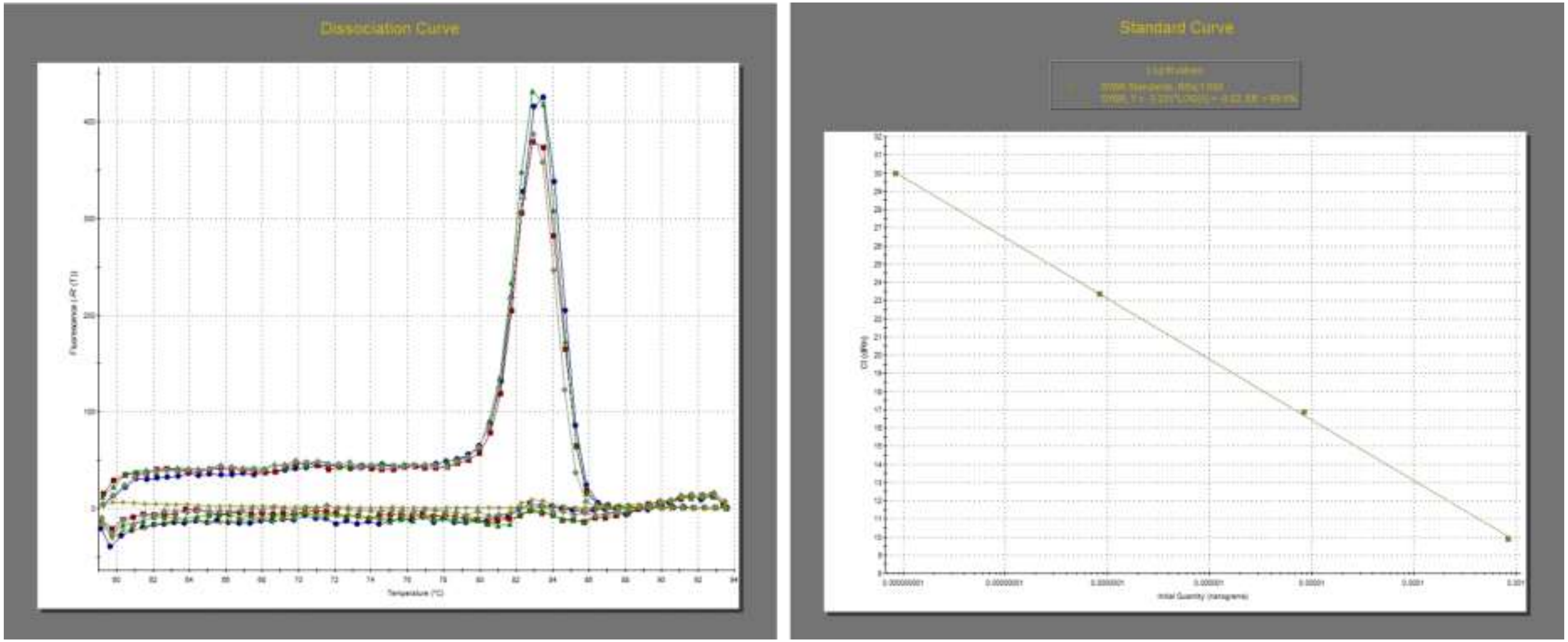


**Supplementary Figure S7.** Dissociation curve and standard curve plots for MdEV1 qPCRs.

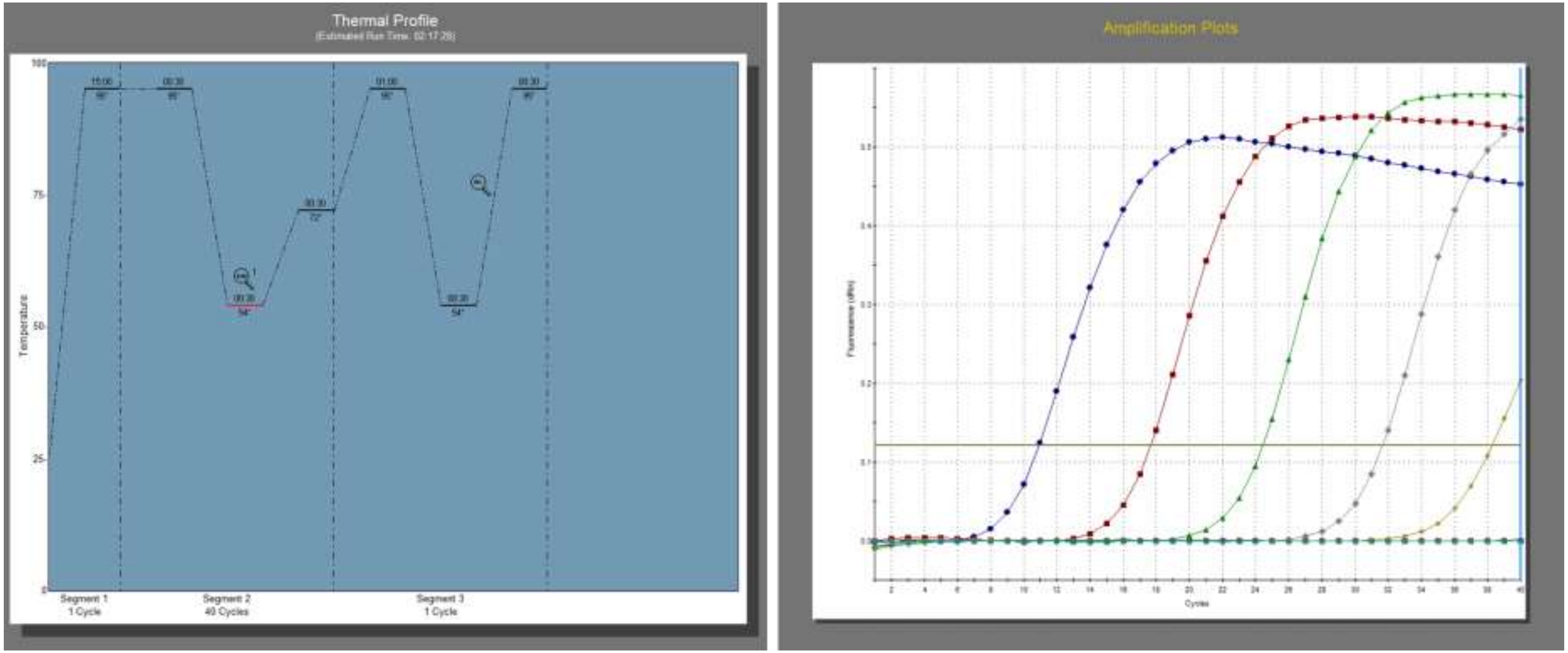


**Supplementary Figure S8.** Thermal profile and amplification plots for MdEV2 qPCRs.

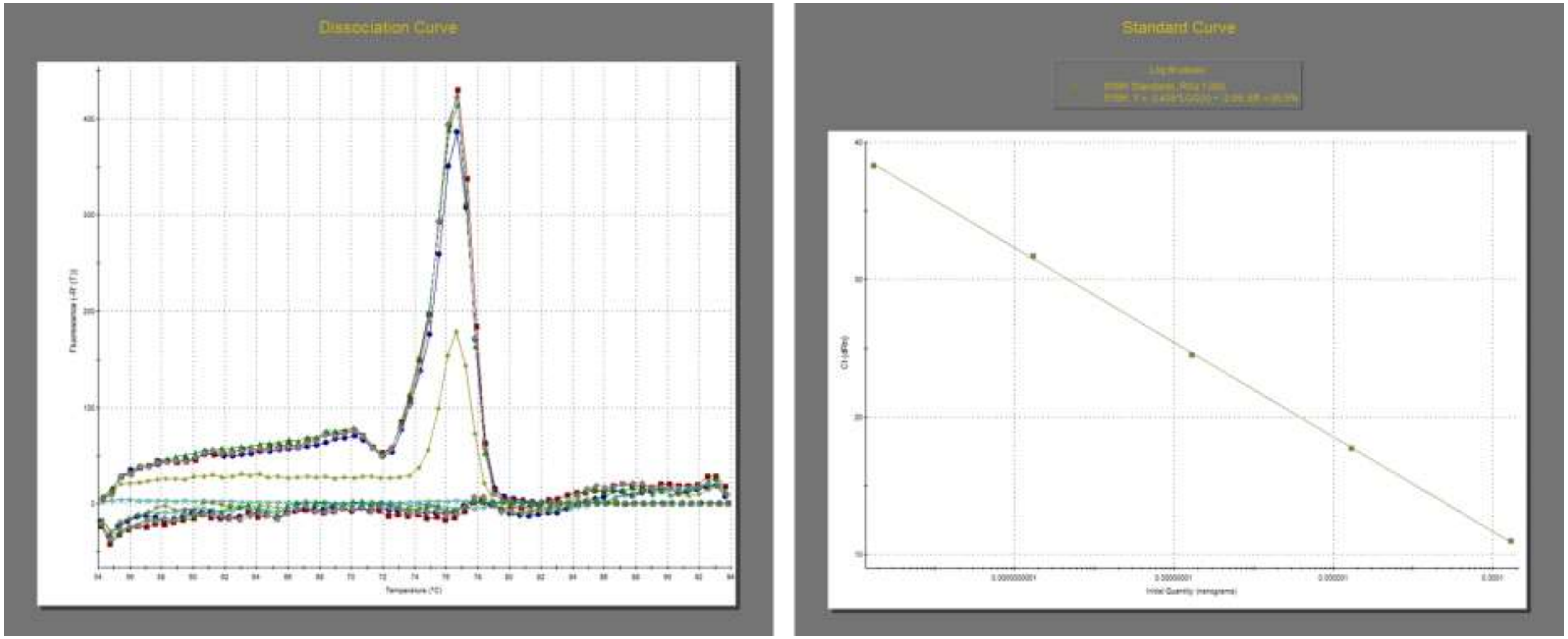


**Supplementary Figure S9.** Dissociation curve and standard curve plots for MdEV2 qPCRs.

**Supplementary Tables**

**Supplementary Table S1.** Number and percentage reads mapped to the housefly, *E. muscae* and virus genomes, and those unmapped from RNAseq of pooled housefly heads.

| Sample | Total reads | Housefly | | *E. muscae* | | Virus | | Unmapped | |
|---|---|---|---|---|---|---|---|---|---|
| | | Number | Percent | Number | Percent | Number | Percent | Number | Percent |
| **CTR_1** | 24,311,249 | 21,690,488 | 89.2 | 20,974 | 0.1 | 2,492 | 0.0 | 2,597,295 | 10.7 |
| **CTR_2** | 24,805,968 | 22,025,933 | 88.8 | 18,401 | 0.1 | 2,435 | 0.0 | 2,759,199 | 11.1 |
| **CTR_3** | 19,992,504 | 17,774,978 | 88.9 | 16,756 | 0.1 | 2,115 | 0.0 | 2,198,655 | 11.0 |
| **CTR_4** | 21,406,316 | 19,387,926 | 90.6 | 22,132 | 0.1 | 2,951 | 0.0 | 1,993,307 | 9.3 |
| **CTR_5** | 23,442,219 | 20,478,604 | 87.4 | 23,314 | 0.1 | 2,098 | 0.0 | 2,938,203 | 12.5 |
| **IC_1** | 24,982,629 | 7,187,959 | 28.8 | 40,115 | 0.2 | 16,824,973 | 67.3 | 929,582 | 3.7 |
| **IC_2** | 24,301,519 | 6,158,356 | 25.3 | 43,816 | 0.2 | 17,282,060 | 71.1 | 817,287 | 3.4 |
| **IC_3** | 21,812,738 | 7,526,252 | 34.5 | 44,431 | 0.2 | 13,299,498 | 61.0 | 942,557 | 4.3 |

| | | | | | | | | | |
|---|---|---|---|---|---|---|---|---|---|
| **IC_4** | 27,346,298 | 4,944,680 | 18.1 | 43,862 | 0.2 | 21,791,293 | 79.7 | 566,463 | 2.1 |
| **IC_5** | 23,522,555 | 4,423,429 | 18.8 | 61,536 | 0.3 | 18,474,066 | 78.5 | 563,524 | 2.4 |
| **SM_1** | 25,006,754 | 6,802,776 | 27.2 | 82,581 | 0.3 | 17,201,803 | 68.8 | 919,594 | 3.7 |
| **SM_2** | 23,452,973 | 3,734,368 | 15.9 | 43,720 | 0.2 | 19,240,624 | 82.0 | 434,261 | 1.9 |
| **SM_3** | 24,724,747 | 3,714,855 | 15.0 | 47,711 | 0.2 | 20,593,164 | 83.3 | 369,017 | 1.5 |
| **SM_4** | 20,207,793 | 5,527,557 | 27.4 | 66,797 | 0.3 | 13,872,178 | 68.6 | 741,261 | 3.7 |
| **SM_5** | 26,376,959 | 5,196,055 | 19.7 | 42,586 | 0.2 | 20,581,488 | 78.0 | 556,830 | 2.1 |
| **PB_1** | 26,218,384 | 1,972,299 | 7.5 | 42,721 | 0.2 | 23,945,466 | 91.3 | 257,898 | 1.0 |
| **PB_2** | 23,956,041 | 5,872,865 | 24.5 | 140,633 | 0.6 | 17,201,579 | 71.8 | 740,964 | 3.1 |
| **PB_3** | 22,736,630 | 4,239,830 | 18.6 | 168,534 | 0.7 | 17,808,565 | 78.3 | 519,701 | 2.3 |
| **PB_4** | 22,662,473 | 3,077,816 | 13.6 | 59,090 | 0.3 | 19,131,764 | 84.4 | 393,803 | 1.7 |
| **PB_5** | 21,133,323 | 1,193,615 | 5.6 | 129,010 | 0.6 | 19,650,097 | 93.0 | 160,601 | 0.8 |
| **WG_1** | 25,530,710 | 2,823,234 | 11.1 | 88,525 | 0.3 | 22,372,016 | 87.6 | 246,935 | 1.0 |

| | | | | | | | | | |
|---|---|---|---|---|---|---|---|---|---|
| **WG_2** | 23,906,841 | 2,681,540 | 11.2 | 97,541 | 0.4 | 20,692,254 | 86.6 | 435,506 | 1.8 |
| **WG_3** | 23,430,536 | 2,746,669 | 11.7 | 76,396 | 0.3 | 20,221,737 | 86.3 | 385,734 | 1.6 |
| **WG_4** | 19,731,281 | 796,181 | 4.0 | 27,460 | 0.1 | 18,756,656 | 95.1 | 150,984 | 0.8 |
| **WG_5** | 25,089,160 | 3,112,229 | 12.4 | 77,693 | 0.3 | 21,510,607 | 85.7 | 388,631 | 1.5 |

**Supplementary Table S2.** Number of raw RNAseq reads that mapped to the different *E. muscae* virus genomes from pooled housefly heads.

| **Sample** | **KVL21-41 EV** | **KVL21-43 EV** | **KVL21-42 EV1** | **KVL21-42 EV2** | **KVL21-40 EV** | ***Delia radicum E. muscae* EV** | **Twyford Virus** | ***E. muscae* Berkeley EV** |
|---|---|---|---|---|---|---|---|---|
| **CTR_1** | 0 | 0 | 0 | 0 | 2,492 | 0 | 0 | 0 |
| **CTR_2** | 0 | 0 | 0 | 0 | 2,435 | 0 | 0 | 0 |
| **CTR_3** | 0 | 0 | 0 | 0 | 2,115 | 0 | 0 | 0 |

| | | | | | | | | |
|---|---|---|---|---|---|---|---|---|
| **CTR_4** | 0 | 0 | 0 | 0 | 2,951 | 0 | 0 | 0 |
| **CTR_5** | 0 | 0 | 0 | 0 | 2,098 | 0 | 0 | 0 |
| **IC_1** | 0 | 1 | 1 | 1 | 16,825,000 | 2 | 7 | 9 |
| **IC_2** | 0 | 1 | 1 | 1 | 17,282,000 | 2 | 4 | 7 |
| **IC_3** | 0 | 2 | 2 | 2 | 13,299,500 | 0 | 4 | 11 |
| **IC_4** | 0 | 2 | 2 | 2 | 21,791,300 | 6 | 6 | 4 |
| **IC_5** | 0 | 2 | 2 | 2 | 18,474,000 | 5 | 8 | 3 |
| **SM_1** | 0 | 1 | 1 | 1 | 17,201,800 | 5 | 10 | 7 |
| **SM_2** | 0 | 1 | 1 | 1 | 19,240,600 | 7 | 10 | 3 |
| **SM_3** | 0 | 1 | 1 | 1 | 20,593,100 | 4 | 4 | 6 |
| **SM_4** | 0 | 1 | 1 | 1 | 13,872,200 | 3 | 2 | 4 |
| **SM_5** | 0 | 2 | 2 | 2 | 20,581,500 | 2 | 10 | 1 |
| **PB_1** | 0 | 2 | 2 | 2 | 23,945,400 | 1 | 7 | 2 |

| | | | | | | | | |
|---|---|---|---|---|---|---|---|---|
| **PB_2** | 0 | 2 | 2 | 2 | 17,201,600 | 3 | 10 | 5 |
| **PB_3** | 0 | 2 | 2 | 2 | 17,808,500 | 3 | 4 | 2 |
| **PB_4** | 0 | 1 | 1 | 1 | 19,131,700 | 5 | 9 | 3 |
| **PB_5** | 0 | 1 | 1 | 1 | 19,650,100 | 6 | 6 | 4 |
| **WG_1** | 0 | 1 | 1 | 1 | 22,372,000 | 5 | 16 | 6 |
| **WG_2** | 0 | 2 | 2 | 2 | 20,692,200 | 3 | 6 | 8 |
| **WG_3** | 0 | 1 | 1 | 1 | 20,221,700 | 4 | 6 | 6 |
| **WG_4** | 0 | 1 | 1 | 1 | 18,756,600 | 5 | 8 | 6 |
| **WG_5** | 0 | 1 | 1 | 1 | 21,510,600 | 2 | 8 | 7 |

**Supplementary Table S3.** The predicted protein-protein interactions between *Entomophthora muscae* candidate secreted proteins identified to be putatively involved in summiting behaviour and the host proteome.

| E. muscae gene | Host transcript | PPI score | Product description |
|---|---|---|---|
| **HBVY01004929** | MDOA012501-RA | 0.518149 | phosphoglycerate mutase 2 [Source:RefSeq gene name;Acc:LOC101891322] |
| **HBVY01007884** | MDOA006559-RA | 0.758677 | formin-A [Source:RefSeq gene name;Acc:LOC101892894] |
| **HBVY01016886** | MDOA013345-RB | 0.85982 | tyrosine-protein phosphatase 99A-like, partial [Source:RefSeq gene name;Acc:LOC101892878] |
| **HBVY01018869** | MDOA010189-RB | 0.623106 | T-complex protein 1 subunit eta [Source:RefSeq gene name;Acc:LOC101887278] |
| | MDOA007621-RA | 0.945118 | T-complex protein 1 subunit beta [Source:RefSeq gene name;Acc:LOC101887429] |
| | MDOA013636-RA | 0.877334 | T-complex protein 1 subunit theta [Source:RefSeq gene name;Acc:LOC101887947] |
| | MDOA006225-RA | 0.973122 | T-complex protein 1 subunit delta [Source:RefSeq gene name;Acc:LOC101889634] |
| | MDOA003290-RA | 0.728611 | T-complex protein 1 subunit alpha [Source:RefSeq gene name;Acc:LOC101891344] |
| | MDOA004575-RA | 0.625952 | T-complex protein 1 subunit gamma [Source:RefSeq gene name;Acc:LOC101892096] |
| | MDOA007155-RA | 0.948308 | 60 kDa heat shock protein, mitochondrial [Source:RefSeq gene name;Acc:LOC101896810] |

| | | | |
|---|---|---|---|
| **HBVY01018869** | MDOA013063-RA | 0.596097 | T-complex protein 1 subunit epsilon [Source:RefSeq gene name;Acc:LOC101897099] |
| | MDOA002569-RA | 0.942979 | T-complex protein 1 subunit zeta [Source:RefSeq gene name;Acc:LOC101899826] |
| **HBVY01021106** | MDOA012295-RA | 0.874739 | cytochrome b5 [Source:RefSeq gene name;Acc:LOC101895524] |
| | MDOA012295-RB | 0.937507 | cytochrome b5 [Source:RefSeq gene name;Acc:LOC101895524] |
| | MDOA013422-RA | 0.502508 | cytochrome b5 [Source:RefSeq gene name;Acc:LOC101898889] |
| | MDOA003707-RE | 0.513154 | synaptotagmin 1 [Source:RefSeq gene name;Acc:LOC101899869] |
| | MDOA003707-RF | 0.772001 | synaptotagmin 1 [Source:RefSeq gene name;Acc:LOC101899869] |
| | MDOA016884-RA | 0.887879 | Down syndrome cell adhesion molecule-like protein Dscam2, partial [Source:RefSeq gene name;Acc:LOC105261444] |
| | MDOA016553-RA | 0.69008 | Down syndrome cell adhesion molecule-like protein Dscam2 [Source:RefSeq gene name;Acc:LOC105262089] |
| | MDOA015916-RA | 0.859618 | tyrosine-protein phosphatase 99A-like, partial [Source:RefSeq gene name;Acc:LOC105262409] |
| **HBVY01016605** | MDOA009434-RA | 0.816993 | Alpha-amylase [Source:UniProtKB/TrEMBL;Acc:A0A1I8MXJ4] |

| | | | |
|---|---|---|---|
| HBVY01016605 | MDOA004496-RA | 0.919039 | Uncharacterized conserved protein [Source:Projected from Glossina morsitans (GMOY007791) UniProtKB/TrEMBL;Acc:D3TPB8] |
| | MDOA013907-RA | 0.687378 | monocarboxylate transporter 4 [Source:RefSeq gene name;Acc:LOC101887574] |
| | MDOA003667-RC | 0.600151 | facilitated trehalose transporter Tret1-like [Source:RefSeq gene name;Acc:LOC101888600] |
| | MDOA011477-RB | 0.761694 | mucin-2-like [Source:RefSeq gene name;Acc:LOC101888831] |
| | MDOA011477-RC | 0.723554 | mucin-2-like [Source:RefSeq gene name;Acc:LOC101888831] |
| | MDOA011477-RD | 0.742873 | mucin-2-like [Source:RefSeq gene name;Acc:LOC101888831] |
| | MDOA003653-RB | 0.815653 | facilitated trehalose transporter Tret1-like [Source:RefSeq gene name;Acc:LOC101889072] |
| | MDOA000729-RA | 0.67936 | 39S ribosomal protein L4, mitochondrial [Source:RefSeq gene name;Acc:LOC101889150] |
| | MDOA011325-RA | 0.715403 | putative inorganic phosphate cotransporter [Source:RefSeq gene name;Acc:LOC101889157] |
| | MDOA009525-RA | 0.775519 | facilitated trehalose transporter Tret1-like [Source:RefSeq gene name;Acc:LOC101889246] |

| | | | |
|---|---|---|---|
| **HBVY01016605** | MDOA001725-RA | 0.646471 | glycogenin-1 [Source:RefSeq gene name;Acc:LOC101889378] |
| | MDOA009720-RC | 0.678649 | facilitated trehalose transporter Tret1-2 homolog [Source:RefSeq gene name;Acc:LOC101889471] |
| | MDOA000363-RA | 0.769252 | 39S ribosomal protein L2, mitochondrial [Source:RefSeq gene name;Acc:LOC101889650] |
| | MDOA007397-RB | 0.838243 | cytochrome b-c1 complex subunit Rieske, mitochondrial [Source:RefSeq gene name;Acc:LOC101889907] |
| | MDOA012019-RA | 0.871495 | protein FAM114A2 [Source:RefSeq gene name;Acc:LOC101890027] |
| | MDOA004678-RA | 0.509999 | mannosyl-oligosaccharide alpha-1,2-mannosidase isoform A-like [Source:RefSeq gene name;Acc:LOC101890227] |
| | MDOA006568-RA | 0.756017 | pre-mRNA 3'-end-processing factor FIP1-like [Source:RefSeq gene name;Acc:LOC101890493] |
| | MDOA003803-RB | 0.753578 | transient receptor potential protein [Source:RefSeq gene name;Acc:LOC101890597] |
| | MDOA012230-RB | 0.748645 | dopamine D2-like receptor [Source:RefSeq gene name;Acc:LOC101890825] |
| | MDOA007223-RF | 0.681135 | dnaJ homolog subfamily B member 6 [Source:RefSeq gene name;Acc:LOC101890834] |

| | | | |
|---|---|---|---|
| **HBVY01016605** | MDOA012350-RA | 0.802633 | 60S acidic ribosomal protein P0 [Source:RefSeq gene name;Acc:LOC101890897] |
| | MDOA011787-RA | 0.915432 | peroxisomal membrane protein PEX14 [Source:RefSeq gene name;Acc:LOC101890933] |
| | MDOA013908-RA | 0.962855 | cytochrome c oxidase subunit 5A, mitochondrial [Source:RefSeq gene name;Acc:LOC101891283] |
| | MDOA010794-RB | 0.663255 | chromosomal protein D1-like [Source:RefSeq gene name;Acc:LOC101891363] |
| | MDOA014508-RB | 0.657656 | protein cereblon homolog [Source:RefSeq gene name;Acc:LOC101891487] |
| | MDOA010371-RA | 0.73224 | transcription factor grauzone-like [Source:RefSeq gene name;Acc:LOC101891521] |
| | MDOA006298-RA | 0.579167 | ADP-ribosylation factor-like protein 16 [Source:RefSeq gene name;Acc:LOC101892061] |
| | MDOA000339-RB | 0.536672 | calnexin [Source:RefSeq gene name;Acc:LOC101892312] |
| | MDOA001829-RA | 0.546562 | homeobox protein 13 [Source:RefSeq gene name;Acc:LOC101892375] |
| | MDOA012492-RC | 0.85504 | threonine--tRNA ligase, cytoplasmic [Source:RefSeq gene name;Acc:LOC101892549] |
| | MDOA010022-RB | 0.816624 | cytosolic purine 5'-nucleotidase [Source:RefSeq gene name;Acc:LOC101893244] |

| | | | |
|---|---|---|---|
| | MDOA010022-RC | 0.732572 | cytosolic purine 5'-nucleotidase [Source:RefSeq gene name;Acc:LOC101893244] |
| | MDOA010022-RD | 0.791216 | cytosolic purine 5'-nucleotidase [Source:RefSeq gene name;Acc:LOC101893244] |
| | MDOA010022-RF | 0.816624 | cytosolic purine 5'-nucleotidase [Source:RefSeq gene name;Acc:LOC101893244] |
| | MDOA010022-RG | 0.816624 | cytosolic purine 5'-nucleotidase [Source:RefSeq gene name;Acc:LOC101893244] |
| | MDOA010022-RH | 0.765842 | cytosolic purine 5'-nucleotidase [Source:RefSeq gene name;Acc:LOC101893244] |
| | MDOA013296-RB | 0.665605 | basic proline-rich protein-like [Source:RefSeq gene name;Acc:LOC101893432] |
| **HBVY01016605** | MDOA002470-RB | 0.69831 | bromodomain-containing protein DDB_G0270170 [Source:RefSeq gene name;Acc:LOC101893453] |
| | MDOA013315-RA | 0.770446 | cytochrome c oxidase subunit 5B, mitochondrial [Source:RefSeq gene name;Acc:LOC101893513] |
| | MDOA004042-RC | 0.765619 | G1/S-specific cyclin-E [Source:RefSeq gene name;Acc:LOC101893514] |
| | MDOA002698-RB | 0.850923 | zinc finger protein 391 [Source:RefSeq gene name;Acc:LOC101894057] |
| | MDOA004877-RA | 0.829847 | putative inorganic phosphate cotransporter [Source:RefSeq gene name;Acc:LOC101894074] |

| | | | |
|---|---|---|---|
| | MDOA003370-RA | 0.662049 | cGMP-dependent protein kinase, isozyme 2 forms cD4/T1/T3A/T3B-like [Source:RefSeq gene name;Acc:LOC101894230] |
| | MDOA004918-RB | 0.67721 | LOW QUALITY PROTEIN: sialin [Source:RefSeq gene name;Acc:LOC101894398] |
| | MDOA004491-RA | 0.693244 | calcium uptake protein 3, mitochondrial [Source:RefSeq gene name;Acc:LOC101894408] |
| | MDOA010921-RA | 0.526172 | putative inorganic phosphate cotransporter [Source:RefSeq gene name;Acc:LOC101895176] |
| **HBVY01016605** | MDOA001227-RA | 0.764192 | sialin [Source:RefSeq gene name;Acc:LOC101895374] |
| | MDOA010897-RA | 0.72635 | putative inorganic phosphate cotransporter [Source:RefSeq gene name;Acc:LOC101895520] |
| | MDOA001406-RA | 0.96009 | cytochrome c oxidase assembly protein COX11, mitochondrial [Source:RefSeq gene name;Acc:LOC101895527] |
| | MDOA012826-RA | 0.814569 | arginine/serine-rich protein PNISR [Source:RefSeq gene name;Acc:LOC101895548] |
| | MDOA010381-RA | 0.610736 | putative inorganic phosphate cotransporter [Source:RefSeq gene name;Acc:LOC101895644] |

| | | | |
|---|---|---|---|
| | MDOA004977-RB | 0.695501 | putative inorganic phosphate cotransporter [Source:RefSeq gene name;Acc:LOC101895696] |
| | MDOA007984-RA | 0.847468 | 14-3-3 protein zeta [Source:RefSeq gene name;Acc:LOC101895723] |
| | MDOA007984-RB | 0.8247 | 14-3-3 protein zeta [Source:RefSeq gene name;Acc:LOC101895723] |
| | MDOA007984-RC | 0.847468 | 14-3-3 protein zeta [Source:RefSeq gene name;Acc:LOC101895723] |
| | MDOA007984-RD | 0.847468 | 14-3-3 protein zeta [Source:RefSeq gene name;Acc:LOC101895723] |
| **HBVY01016605** | MDOA014956-RA | 0.737774 | leucine-rich PPR motif-containing protein, mitochondrial [Source:RefSeq gene name;Acc:LOC101895782] |
| | MDOA012840-RA | 0.8646 | putative inorganic phosphate cotransporter [Source:RefSeq gene name;Acc:LOC101895873] |
| | MDOA006173-RB | 0.852796 | putative inorganic phosphate cotransporter [Source:RefSeq gene name;Acc:LOC101896045] |
| | MDOA001214-RA | 0.750218 | BUD13 homolog [Source:RefSeq gene name;Acc:LOC101896051] |
| | MDOA008204-RA | 0.506244 | signal recognition particle 54 kDa protein [Source:RefSeq gene name;Acc:LOC101896252] |

| | | | |
|---|---|---|---|
| | MDOA000012-RA | 0.669095 | caspase Nc [Source:RefSeq gene name;Acc:LOC101896806] |
| | MDOA005140-RB | 0.940877 | hexaprenyldihydroxybenzoate methyltransferase, mitochondrial-like [Source:RefSeq gene name;Acc:LOC101896946] |
| | MDOA005140-RC | 0.81828 | hexaprenyldihydroxybenzoate methyltransferase, mitochondrial-like [Source:RefSeq gene name;Acc:LOC101896946] |
| | MDOA005140-RD | 0.937942 | hexaprenyldihydroxybenzoate methyltransferase, mitochondrial-like [Source:RefSeq gene name;Acc:LOC101896946] |
| **HBVY01016605** | MDOA005527-RA | 0.604582 | iron-sulfur cluster assembly 1 homolog, mitochondrial [Source:RefSeq gene name;Acc:LOC101897302] |
| | MDOA009799-RA | 0.889286 | enhancer of filamentation 1-like [Source:RefSeq gene name;Acc:LOC101897309] |
| | MDOA002251-RA | 0.582174 | 60S ribosomal protein L23 [Source:RefSeq gene name;Acc:LOC101897495] |
| | MDOA001185-RA | 0.671342 | solute carrier family 12 member 6 [Source:RefSeq gene name;Acc:LOC101897946] |
| | MDOA001185-RB | 0.671341 | solute carrier family 12 member 6 [Source:RefSeq gene name;Acc:LOC101897946] |
| | MDOA001185-RC | 0.535582 | solute carrier family 12 member 6 [Source:RefSeq gene name;Acc:LOC101897946] |
| | MDOA001185-RD | 0.860931 | solute carrier family 12 member 6 [Source:RefSeq gene name;Acc:LOC101897946] |

| | | | |
|---|---|---|---|
| HBVY01016605 | MDOA001185-RE | 0.861718 | solute carrier family 12 member 6 [Source:RefSeq gene name;Acc:LOC101897946] |
| | MDOA001185-RF | 0.726159 | solute carrier family 12 member 6 [Source:RefSeq gene name;Acc:LOC101897946] |
| | MDOA014844-RA | 0.749874 | facilitated trehalose transporter Tret1-2 homolog [Source:RefSeq gene name;Acc:LOC101898182] |
| | MDOA013703-RA | 0.56963 | 28S ribosomal protein S15, mitochondrial [Source:RefSeq gene name;Acc:LOC101898935] |
| | MDOA006775-RA | 0.709032 | ribosomal RNA-processing protein 17 [Source:RefSeq gene name;Acc:LOC101899029] |
| | MDOA004583-RA | 0.546332 | endoplasmin [Source:RefSeq gene name;Acc:LOC101899169] |
| | MDOA006094-RB | 0.979234 | NADH-ubiquinone oxidoreductase 75 kDa subunit, mitochondrial [Source:RefSeq gene name;Acc:LOC101899436] |
| | MDOA014443-RA | 0.88668 | peroxiredoxin-5, mitochondrial [Source:RefSeq gene name;Acc:LOC101899846] |
| | MDOA015392-RA | 0.761091 | lipoamide acyltransferase component of branched-chain alpha-keto acid dehydrogenase complex, mitochondrial [Source:RefSeq gene name;Acc:LOC101900261] |
| HBVY01016605 | MDOA015392-RB | 0.580436 | lipoamide acyltransferase component of branched-chain alpha-keto acid dehydrogenase complex, mitochondrial [Source:RefSeq gene name;Acc:LOC101900261] |

| | | | |
|---|---|---|---|
| | MDOA006600-RA | 0.946735 | FAD-linked sulfhydryl oxidase ALR [Source:RefSeq gene name;Acc:LOC101900794] |
| | MDOA013348-RA | 0.687735 | E3 ubiquitin-protein ligase MARCH6 [Source:RefSeq gene name;Acc:LOC101901242] |
| | MDOA005693-RA | 0.949397 | sialin [Source:RefSeq gene name;Acc:LOC101901262] |
| | MDOA005693-RB | 0.949397 | sialin [Source:RefSeq gene name;Acc:LOC101901262] |
| | MDOA001400-RI | 0.778344 | sodium/calcium exchanger 3-like [Source:RefSeq gene name;Acc:LOC101901283] |
| | MDOA008256-RB | 0.515534 | solute carrier family 52, riboflavin transporter, member 3-A [Source:RefSeq gene name;Acc:LOC101901351] |
| | MDOA008256-RD | 0.515534 | solute carrier family 52, riboflavin transporter, member 3-A [Source:RefSeq gene name;Acc:LOC101901351] |
| | MDOA011886-RA | 0.802254 | 40S ribosomal protein S21 [Source:RefSeq gene name;Acc:LOC101901468] |
| | MDOA010526-RA | 0.907972 | mucin-2-like, partial [Source:RefSeq gene name;Acc:LOC101901695] |
| | MDOA016167-RA | 0.975238 | facilitated trehalose transporter Tret1 [Source:RefSeq gene name;Acc:LOC105261430] |
| **HBVY01016605** | MDOA016733-RA | 0.696126 | putative inorganic phosphate cotransporter [Source:RefSeq gene name;Acc:LOC105261617] |

| | | | |
|---|---|---|---|
| | MDOA016713-RB | 0.820133 | nucleolar and coiled-body phosphoprotein 1 [Source:RefSeq gene name;Acc:LOC105262084] |
| | MDOA016713-RC | 0.811469 | nucleolar and coiled-body phosphoprotein 1 [Source:RefSeq gene name;Acc:LOC105262084] |
| | MDOA016531-RA | 0.549214 | putative inorganic phosphate cotransporter [Source:RefSeq gene name;Acc:LOC105262542] |
| | MDOA011775-RC | 0.798061 | Leucine Rich repeat protein [Source:UniProtKB/TrEMBL;Acc:T1PMW8] |
| | MDOA000525-RA | 0.6758 | unspecified product |
| | MDOA014249-RA | 0.682976 | unspecified product |
| | MDOA002128-RA | 0.682976 | unspecified product |
| | MDOA010234-RA | 0.532135 | unspecified product |
| | MDOA013469-RA | 0.619054 | unspecified product |
| **HBVY01016605** | MDOA012959-RA | 0.885931 | unspecified product |
| | MDOA011633-RA | 0.638556 | unspecified product |

**Supplementary Table S4.** The predicted protein-protein interactions between the partial KVL21-40 *E. muscae* virus protease protein and the host proteome.

| Host transcript | PPI score | Product description |
|---|---|---|
| MDOA013387-RA | 0.629093945 | ecdysone receptor [Source:RefSeq gene name;Acc:LOC101888839] |
| MDOA003144-RA | 0.849262059 | protein claret segregational [Source:RefSeq gene name;Acc:LOC101891358] |
| MDOA009609-RA | 0.89308697 | transcription factor HNF-4 homolog [Source:RefSeq gene name;Acc:LOC101891665] |
| MDOA003867-RA | 0.500161409 | kinesin-like protein KIF3A [Source:RefSeq gene name;Acc:LOC101898167] |
| MDOA013977-RB | 0.573267698 | probable nuclear hormone receptor HR38 [Source:RefSeq gene name;Acc:LOC101898949] |
| MDOA010970-RB | 0.781938434 | kinesin-like protein Nod [Source:RefSeq gene name;Acc:LOC101900234] |
| MDOA007676-RC | 0.503721654 | chromosome-associated kinesin KIF4 [Source:RefSeq gene name;Acc:LOC101900623] |
